# Full genome *Nobecovirus* sequences from Malagasy fruit bats define a unique evolutionary history for this coronavirus clade

**DOI:** 10.1101/2021.09.29.462406

**Authors:** Gwenddolen Kettenburg, Amy Kistler, Hafaliana Christian Ranaivoson, Vida Ahyong, Angelo Andrianiaina, Santino Andry, Joseph L. DeRisi, Anecia Gentles, Vololoniaina Raharinosy, Tsiry Hasina Randriambolamanantsoa, Ny Anjara Fifi Ravelomanantsoa, Cristina M. Tato, Philippe Dussart, Jean-Michel Heraud, Cara E. Brook

## Abstract

Bats are natural reservoirs for both *Alpha*- and *Betacoronaviruses* and the hypothesized original hosts of five of seven known zoonotic coronaviruses. To date, the vast majority of bat coronavirus research has been concentrated in Asia, though coronaviruses are globally distributed; indeed, SARS-CoV and SARS-CoV-2-related *Betacoronaviruses* in the subgenus *Sarbecovirus* have been identified circulating in *Rhinolophid* bats in both Africa and Europe, despite the relative dearth of surveillance in these regions. As part of a long-term study examining the dynamics of potentially zoonotic viruses in three species of endemic Madagascar fruit bat (*Pteropus rufus, Eidolon dupreanum, Rousettus madagascariensis*), we carried out metagenomic Next Generation Sequencing (mNGS) on urine, throat, and fecal samples obtained from wild-caught individuals. We report detection of RNA derived from *Betacoronavirus* subgenus *Nobecovirus* in fecal samples from all three species and describe full genome sequences of novel *Nobecoviruses* in *P. rufus* and *R. madagascariensis*. Phylogenetic analysis indicates the existence of five distinct *Nobecovirus* clades, one of which is defined by the highly divergent sequence reported here from *P. rufus* bats. Madagascar *Nobecoviruses* derived from *P. rufus* and *R. madagascariensis* demonstrate, respectively, Asian and African phylogeographic origins, mirroring those of their fruit bat hosts. Bootscan recombination analysis indicates significant selection has taken place in the spike, nucleocapsid, and NS7 accessory protein regions of the genome for viruses derived from both bat hosts. Madagascar offers a unique phylogeographic nexus of bats and viruses with both Asian and African phylogeographic origins, providing opportunities for unprecedented mixing of viral groups and, potentially, recombination. As fruit bats are handled and consumed widely across Madagascar for subsistence, understanding the landscape of potentially zoonotic coronavirus circulation is essential for mitigation of future zoonotic threats.

## Introduction

In the past 20 years, bat-derived coronaviruses SARS-CoV, MERS-CoV, and SARS-CoV-2 have been responsible for two deadly epidemics and the ongoing COVID-19 pandemic (1–4). These coronaviruses (CoVs) are members of the *Betacoronavirus* genus, which, along with genus *Alphacoronavirus*, are primarily associated with bat hosts (1–4); the remaining CoV genera, *Gammacoronavirus and Deltacoronavirus*, are typically hosted by birds (5). The *Betacoronavirus* group can be further broken down into five subgenera: *Sarbecovirus* (hosted by bats in family Rhinolophidae (6,7)), *Merbecovirus* (hosted by bats in family Vespertilionidae (8– 10)), *Nobecovirus* (hosted by bats in family Pteropodidae (11–13)), and *Hibecovirus* (hosted by bats in family Hipposideridae (14–16)). The fifth *Betacoronavirus* subgenus, *Embecovirus*, is primarily associated with rodents and bovids, though a few bat hosts have been documented (17,18). Since the emergence of SARS-CoV in 2002, there has been increasing interest in surveying potential hosts of coronaviruses and contributing new virus sequences to public databases, with most effort focused on sampling bats from Asia (19–27), the continent of origin for both the SARS-CoV epidemic and the SARS-CoV-2 pandemic. Recently, more concerted efforts have arisen to survey the landscape of bat-borne coronaviruses in other regions of the world, including Africa and Europe (11,12,28–32).

The family *Coronaviridae* is considered one of the most likely viral taxa to switch host species (33,34) in part because many CoVs utilize well-conserved cell surface receptors presented on a wide variety of mammalian host cells. The zoonotic *Sarbecoviruses*, SARS-CoV and SARS-CoV-2, for example, use the human cell surface receptor Angiotensin-converting enzyme 2 (ACE2) to gain entry into human cells (35,36), while many *Merbecoviruses* interact with the well-conserved vertebrate host cell receptor dipeptidyl peptidase 4 (DPP4) (37). As only a fraction of the *Alpha-* and *Betacoronavirus* diversity projected to circulate in wild bat hosts has been already described (38), it is possible that many CoVs capable of zoonotic emergence remain uncharacterized. Because CoVs are known to recombine with other CoVs, or more rarely, with other viral groups (39–44), there is additional concern that naturally-circulating CoVs presently unable to infect humans may acquire this ability in the future. Several factors, which have been reviewed at length elsewhere (4,33,45), contribute to the propensity for CoV recombination, including the large CoV genome size supported by a unique proofreading mechanism in the RNA-dependent RNA polymerase (RdRp) (46–49), as well as a ‘copy choice’ template switching mechanism of RNA replication whereby RdRp physically detaches from one RNA template during replication and reattaches to an adjacent template, thus facilitating recombination in cases where multiple viruses may be coinfecting the same cell (50).

Madagascar is an island country in southeastern Sub-Saharan Africa, located in the Indian Ocean, ∼400 km off the coast from Mozambique. Madagascar has been isolated from the African continent for over 170 million years and all surrounding landmasses for over 80 million years, allowing for the evolution of a unique and highly endemic floral and faunal assemblage (51). The country is home to 51 species of bat (52), two-thirds of which are endemic and boast long evolutionary divergence times with sister species on both the African and Asian continents (53– 55). A growing body of work has characterized the landscape of potentially zoonotic viruses in Madagascar bats, identifying evidence of circulating infection (through RNA detection or serology) with henipaviruses, filoviruses, lyssaviruses, and coronaviruses (12,32,56,57). Previously, coronavirus surveillance efforts have identified *Alphacoronavirus* RNA in the Malagasy insectivorous bat, *Mormopterus jugularis*, and *Betacoronavirus* RNA in all three endemic Malagasy fruit bat species, *Pteropus rufus, Eidolon dupreanum*, and *Rousettus madagascariensis* (12,32); this latter *Betacoronavirus* RNA clusters with subgenus *Nobecovirus* (12,32), which has been previously characterized from *Pteropodidae* fruit bats across Asia and in both East and West Africa (27,29,58–61). Though *Nobecoviruses* are not known to be zoonotic, previous research has described widespread circulation of a recombinant *Nobecovirus* throughout Asia, which carries an orthoreovirus p10 gene insertion (27,61,62), highlighting the capacity for this viral subgenus to undertake rapid shifts in genomic organization which could lead to expanded host range. As both *E. dupreanum* and *R. madagascariensis* are known to co-roost with each other, and with several species of insectivorous bat (63), CoV recombination is a distinct concern in the Madagascar system. Though no *Rhinolophus* spp. bats, the typical host for ACE2-using *Sarbecoviruses*, inhabit Madagascar, the island is home to four species of Hipposiderid bat (52), which host the *Sarbecovirus-*adjacent and understudied *Hibecoviruses*, as well as several species of Vespertilionid bat, the most common hosts for the zoonotic *Merbecoviruses*.

Human-bat contact rates are high in some regions of Madagascar, where bats are consumed widely for subsistence and frequently roost in close proximity to human settlements or natural tourist attractions (64–68). Moreover, in addition to the natural CoV diversity described in Malagasy bats, several human coronaviruses are known to circulate widely in the human population in Madagascar, including the common cold-causing *Embecoviruses*, HCoV-OC43 and HCoV-HKU1, and, more recently, the zoonotic *Sarbecovirus*, SARS-CoV-2 (69–71). As spillback of SARS-CoV-2 into wildlife hosts and possible recombination with wildlife viruses remains a global concern (16), characterization of the genetic diversity of bat-borne coronaviruses in Madagascar and elsewhere in Africa is a critical public health priority. Here, we contribute and characterize three full genome sequences of two novel *Nobecoviruses*, derived from *R. madagascariensis* and *P. rufus* hosts. We define five distinct *Nobecovirus* clades in global circulation across Asia and Africa and assess these new *Nobecoviruses* for their past and future capacity for recombination.

## Materials and Methods

### Bat Sampling

As part of a long-term study characterizing the seasonal dynamics of potentially zoonotic viruses in wild fruit bats in Madagascar, monthly captures of Malagasy pteropodid bats were carried out at species-specific roost sites in the Districts of Moramanga and Manjakandriana, Madagascar between 2018 and 2019 (*P. rufus:* Ambakoana roost, -18.513 S, 48.167 E; *E. dupreanum*: Angavobe cave, -18.944 S, 47.949 E; Angavokely cave = -18.933 S, 47.758 E; *R. madagascariensis*: Maromizaha cave, -18.9623 S, 48.4525 E). In brief, bats were captured in nets hung in the tree canopy (*P. rufus*) or over cave mouths (*E. dupreanum, R. madagascariensis)* at dusk (17:00-22:00) and dawn (03:00-07:00), removed from nets, and processed under manual restraint following methods that have been previously described (57,72,73). All animals were identified to species, sex, and age class (juvenile vs. adult), and fecal, throat, and urine swabs were taken from each individual, collected into viral transport medium, and frozen on site in liquid nitrogen. Post-sampling, swabs were transported to -80°C freezers for long-term storage in the Virology Unit at Institut Pasteur de Madagascar.

This study was carried out in strict accordance with research permits obtained from the Madagascar Ministry of Forest and the Environment (permit numbers 019/18, 170/18, 007/19) and under guidelines posted by the American Veterinary Medical Association. All field protocols employed were pre-approved by the UC Berkeley Animal Care and Use Committee (ACUC Protocol # AUP-2017-10-10393), and every effort was made to minimize discomfort to animals.

### RNA Extraction

RNA was extracted from a randomly selected subset of fecal, throat, and urine swab samples in the Virology Unit at the Institut Pasteur de Madagascar, with each sample corresponding to a unique individual from the field dataset. Samples undergoing mNGS corresponded to individuals captured in Feb-Apr, Jul-Sep and December 2018 or in January 2019. Water controls were extracted in conjunction with samples on each unique extraction day. Extractions were conducted using the Zymo Quick DNA/RNA Microprep Plus kit (Zymo Research, Irvine, CA, USA), according to the manufacturer’s instructions and including the step for DNAse digestion. Post-extraction, RNA quality was checked on a nanodrop to ensure that all samples demonstrated 260/280 ratios exceeding 2 and revealed quantifiable concentrations. Resulting extractions were stored in freezers at -80°C, then transported on dry ice to the Chan Zuckerberg Biohub (San Francisco, CA, USA) for library preparation and metagenomic Next Generation Sequencing (mNGS).

### Library Preparation and mNGS

Four randomly selected samples from each of three bat species underwent additional quantification using an Invitrogen Qubit 3.0 Fluorometer and the Qubit RNA HS Assay Kit (ThermoFisher Scientific, Carlsbad, CA, USA). After quantification, all total RNA samples, along with water samples from Madagascar extractions, were manually arrayed into 96 well plates to automate high throughput mNGS library preparation. Based on the initial quantitation, a 2uL aliquot from each plated sample was diluted 1:9 on a Bravo liquid handling platform (Agilent, Santa Clara, CA, USA). A 5 *μ*L aliquot from each diluted sample was arrayed into a 384 well plate for input into the mNGS library prep. Samples derived from fecal, throat, and urine swab samples were arrayed on distinct 384 well plates for separate sequencing runs. Additional unrelated total RNA samples (a dilution series of total RNA isolated from cultured HeLa cells) and a set of local lab water samples were included on each 384 well plate to serve as library preparation controls. Input RNA samples in the 384 well plate were transferred to a GeneVac EV-2 (SP Industries, Warminster, PA, USA) to evaporate the samples to enable miniaturized mNGS library preparation with the NEBNext Ultra II RNA Library Prep Kit (New England BioLabs, Beverly, MA, USA). Library preparation was performed per the manufacturer’s instructions, with the following modifications: 25pg of External RNA Controls Consortium Spike-in mix (ERCCS, Thermo-Fisher) was added to each sample prior to RNA fragmentation; the input RNA mixture was fragmented for 8 min at 94°C prior to reverse transcription; and a total of 14 cycles of PCR with dual-indexed TruSeq adapters was applied to amplify the resulting individual libraries. An initial equivolume library pool was generated, and the quality and quantity of that pool was assessed via electrophoresis (High-Sensitivity DNA Kit and Agilent Bioanalyzer; Agilent Technologies, Santa Clara, CA, USA), real-time quantitative polymerase chain reaction (qPCR) (KAPA Library Quantification Kit; Kapa Biosystems, Wilmington, MA, USA), and small-scale sequencing (2 x146bp) on an iSeq platform (Illumina, San Diego, CA, US). Subsequent equimolar pooling of individual libraries from each plate was performed prior to performing large-scale paired-end sequencing (2 × 146 bp) run on the Illumina NovaSeq sequencing system (Illumina, San Diego, CA, USA). The pipeline used to separate the sequencing output of the individual libraries into FASTQ files of 146bp paired-end reads is available on GitHub at https://github.com/czbiohub/utilities.

### IDSeq

Raw reads from Illumina sequencing were host-filtered, quality-filtered, and assembled on the IDseq (v3.10, NR/NT 2019-12-01) platform, a cloud-based, open-source bioinformatics platform designed for microbe detection from metagenomic data (74), using a host background model of “bat” compiled from all publicly available full-length bat genomes in GenBank at the time of sequencing. Samples were deemed “positive” for coronavirus infection if IDseq successfully assembled at least two contigs with an average read depth >2 reads/nucleotide that showed significant nucleotide or protein BLAST alignment(s) (alignment length >100 nt/aa and E-value < 0.00001 for nucleotide BLAST/ bit score >100 for protein BLAST) to any CoV reference present in NCBI NR/NT database (version 12-01-2019). To verify that no positives were missed from IDseq, all non-host contigs assembled in IDseq underwent directed, offline BLASTn and BLASTx (75) against a reference database constructed from all available full-length nucleotide and protein reference sequences for *Alpha-* and *Betacoronavirus* available in NCBI Virus (last access: August 15, 2021). Step-by-step instructions for our offline BLAST protocol can be accessed in our publicly available GitHub repository at: https://github.com/brooklabteam/Mada-Bat-CoV/.

### Genome Annotation and BLAST

Three full genome-length Nobecovirus contigs returned from IDseq (two from *R. madagascariensis* and one from *P. rufus*) were aligned with Nobecovirus homologs from NCBI (see ‘Phylogenetic Analysis’) and annotated in the program Geneious Prime (2020.0.5). We then used NCBI BLAST and BLASTx to query identity of our full length recovered genomes and their respective translated proteins to publicly available sequences in NCBI (75). We queried identity to reference sequences for four previously described *Nobecovirus* strains (accession numbers: MG762674 (*Rousettus* bat coronavirus HKU9), NC_030886 (*Rousettus* bat coronavirus RoBat-CoV GCCDC1), MK211379 (*Rhinolophus affinis* coronavirus BtRt-BetaCoV/GX2018), and NC_048212 (*Eidolon helvum* bat coronavirus), as well as to the top BLAST hit overall. Finally, we aligned representative sequences from each major *Nobecovirus* clade and visually examined the region of p10 orthoreovirus insertion from the RoBat-CoV GCCDC1 lineage in the newly described sequences from Madagascar.

### Phylogenetic Analysis

Contigs returned from IDseq were combined with publicly available coronavirus sequences in NCBI to perform phylogenetic analysis. We carried out four major phylogenetic analyses, building (a) a full-genome *Betacoronavirus* maximum likelihood (ML) phylogeny, (b) a *Betacoronavirus* ML phylogeny corresponding to a conserved 259 bp fragment of the RNA-dependent RNA polymerase (RdRp) gene encapsulated in the CoV Orf1b, (c) four amino acid ML phylogenies derived from translated nucleotides corresponding to the spike (S), envelope (E), matrix (M), and nucleocapsid (N) proteins from a subset of full length genomes, and (d) a *Nobecovirus* Bayesian phylogeny corresponding to a conserved 365 bp subset of the RdRp gene. Detailed methods for the construction of each phylogeny are available at https://github.com/brooklabteam/Mada-Bat-CoV/.

Briefly, our full genome ML phylogeny was comprised of 122 unique NCBI records, corresponding to all available full genome sequences with bat hosts under NCBI taxon IDs, *Betacoronavirus* (694002), unclassified *Betacoronavirus* (696098), *Betacoronavirus* sp. (1928434), unclassified *Coronaviridae* (1986197), or unclassified *Coronavirinae* (2664420) (107 records), in addition to all full genome *Betacoronavirus* (694002) reference sequences with a non-bat host (14 records), plus one *Gammacoronavirus* outgroup (accession number NC_010800). The full genome phylogeny additionally included three full length Madagascar *Nobecovirus* sequences returned from IDseq (two from *R. madagascariensis* and one from *P. rufus*), which are described in this paper for the first time, for a total of 125 unique sequences.

Our *Betacoronavirus* RdRp ML phylogeny consisted of an overlapping subset of a 259 bp fragment in the center of the RdRp gene that has been previously described in Madagascar fruit bats (12) (7 records), in addition to the same RdRp fragment extracted from near-full length *Nobecovirus* sequences on NCBI Virus (17 records) and full length reference sequences for other *Betacoronavirus* subgenera available in NCBI Virus (17 records). This phylogeny also included two NCBI Virus RdRp *Nobecovirus* fragments, in addition to seven Madagascar *Nobecovirus* sequences encompassing the RdRp fragment of interest, which were returned from the assembly in IDseq (four from *R. madagascariensis*, two from *P. rufus*, and one from *E. dupreanum*). Finally, we included the RdRp fragment of our *Gammacoronaviru*s outgroup, for a total of 51 unique sequences.

Our amino acid phylogenies consisted of S, E, M, and N gene extractions from a subset of the same representative set of near-full genome length sequences used in the RdRp analysis: the same 17 full-length *Betacoronavirus* reference sequences, 17 near full-length *Nobecovirus* sequences, and the one *Gammacoronavirus* outgroup, in addition to our three full genome Madagascar sequences derived from *R. madagascariensis* and *P. rufus*. Gene extractions were carried out using annotation tracks reported with each accession number in NCBI or, in cases where annotations were unavailable, genes were manually annotated and extracted in Geneious Prime based on alignment to homologs. After nucleotide extraction, genes were translated prior to alignment.

Finally, our *Nobecovirus* RdRp Bayesian phylogeny consisted of an overlapping subset of a 386 bp fragment of the RdRp gene from a 32-sequence subset of those used in the RdRp ML phylogeny. This included all 13 of 14 possible Madagascar sequences (the one recently described *E. dupreanum* sequence was left out due to its short length), plus all 19 *Nobecovirus* RdRp fragments from NCBI Virus.

After compiling sequences for each phylogenetic analysis, sequence subsets for the full-length, RdRp, and four amino acid phylogenies were aligned in MAFFT v.7 (76,77) using default parameter values. Alignments were checked manually for quality in Geneious Prime, and the RdRp alignments were trimmed a fragment conserved across all sequences in the subset (259 bp for ML phylogeny, 365 for Bayesian phylogeny). All sequence subsets and alignment files are available for public access in our GitHub repository: https://github.com/brooklabteam/Mada-Bat-CoV/.

After quality control, alignments were sent to Modeltest-NG (78) to assess the best fit nucleotide or amino acid substitution model appropriate for the data. All alignments for ML analysis (full genome, 259 bp RdRp fragment, and amino acid protein sequences) were then sent to RAxML-NG (79) to construct the corresponding phylogenetic trees. Following best practices outlined in the RAxML-NG manual, twenty ML inferences were made on each original alignment and bootstrap replicate trees were inferred using Felsenstein’s method (80), with the MRE-based bootstopping test applied after every 50 replicates (81). Bootstrapping was terminated once diagnostic statistics dropped below the threshold value and support values were drawn on the best-scoring tree.

We constructed the Bayesian timetree using the Bayesian Skyline Coalescent (82) model in BEAST2 (83), assuming a constant population prior, and the best fit nucleotide substitution model as indicated by ModelTest-NG. Sampling dates corresponded to collection date as reported in NCBI Virus; in cases where only year was reported, we assumed a collection date of 15-July for the corresponding year. We tested trees using both an uncorrelated exponentially distributed relaxed molecular clock (UCED) and a strict clock. Markov chain Monte Carlo (MCMC) sample chains were run for 1 billion iterations, convergence was checked using TRACER v1.7 (84), and trees were averaged after 10% burn-in using TreeAnnotater v2.6.3 (85) to visualize mean posterior densities at each node.

All resulting phylogenies (both ML and Bayesian) were visualized in R v.4.0.3 for MacIntosh, using the package ‘ggtree’ (86).

### Recombination Analysis

Full length *Nobecovirus* sequences derived from IDseq were analyzed for any signature of past recombination. First, the ORF1a, ORF1b, S, NS3, E, M, N, and NS7 genes from the *P. rufus Nobecovirus* sequence, the longest *R. madagascariensis Nobecovirus* sequence (MIZ240), and two full genome representative sequences from the HKU9 (NC_009021) and *E. helvum* African lineages (NC_048212) were extracted, translated, and concatenated. Concatenated, translated sequences were then aligned, and aligned in MAFFT v.7 (76,77) using default parameter values. *Nobecovirus* sequences corresponding to the RoBat-CoV GCCDC1 (27,61) and BtRt-BetaCoV/GX2018/BtCoV92 (58,59) genotypes were omitted from recombination analyses because inserted genes and/or genetic material upstream from the nucleocapsid in the corresponding genomes interfered with the alignment.

After alignment, genomes were analyzed for amino acid similarity in the program pySimplot (87), using the *P. rufus* and, subsequently, the *R. madagascariensis* genome as query sequences, the HKU9 and *Eidolon helvum* African *Nobecovirus* clades as references, and the corresponding Madagascar sequence as the alternative. Analyses were carried out using a window size of 100aa and a step size of 20aa.

Next, all three full length nucleotide sequences of Madagascar *Nobecovirus* genomes were aligned with grouped full genome sequences corresponding to the two disparate *Nobecovirus* lineages: the HKU9 lineage (EF065514-EF065516, HM211098-HM211100, MG693170, NC_009021, MG762674) and the *E. helvum* African lineage (MG693169, MG693171-MG693172, NC_048212). As before, alignment was conducted in MAFFT v.7 (76,77) using default parameter values.

After alignment, genomes were analyzed for recombination in the program SimPlot (v.3.5.1). Nucleotide similarity plots, were generated using the *P. rufus* and, subsequently, the *R. madagascariensis* genomes as query sequences, the HKU9 and *Eidolon helvum* African *Nobecovirus* clades as references, and the corresponding Madagascar sequence as the alternative. Bootscan analyses were conducted on the same alignment, using the same query and reference inputs. Both nucleotide similarity and Bootscan analyses were carried out using a window size of 200bp and a step size of 20bp.

### Nucleotide Sequence Accession Numbers

All three annotated full-length genome sequences (two from *R. madagascariensis*, one from *P. rufus*), plus four additional RdRp gene fragment sequences (two from *R. madagascariensis*, one from *P. rufus*, and one from *E. dupreanum*) were submitted to NCBI and assigned accession numbers OK020086-OK020089 (RdRp fragments), OK067319-OK067321 (full genomes).

## Summary

### Prevalence of CoV Sequence Detection in Field Specimens

RNA from 285 fecal, 143 throat, and 196 urine swab samples was prepped into libraries and submitted for Illumina sequencing. In 28/285 (9.82%) fecal specimens and in 2/196 (1.00%) urine specimens, at least two contigs with an average read depth > 2 reads/nucleotide, and nucleotide or protein-BLAST alignments to any CoV reference sequence in NCBI were identified via IDseq analysis. Because the prevalence detected in the urine samples was low, it is likely attributable to field contamination with fecal excrement upon urine swab collection, as bats often excrete both substances simultaneously under manual restraint. None of the 143 throat swabs assayed demonstrated evidence of CoV infection.

Prevalence in feces varied slightly across species, with 4/44 (9.1%) *P. rufus* specimens, 16/145 (11.0%) *E. dupreanum* specimens, and 8/96 (8.3%) *R. madagascariensis* specimens sequencing CoV positive. Juveniles demonstrated higher CoV prevalence than adults for *P. rufus* and *E. dupreanum* but not for *R. madagascariensis*. Juvenile vs. adult prevalence was 3/15 (20%) vs. 1/29 (3.5%) for *P. rufus*,5/13 (38.5%) vs. 11/132 (8.3%) for *E. dupreanum*, and 0/13 (0%) vs. 8/83 (9.6%) for *R. madagascariensis* (**Figure 1A**). Prevalence varied seasonally across all three species, peaking coincidentally in adult and juvenile populations for *P. rufus* and *E. dupreanum*, with the highest prevalence for all three species observed during the wet season months of February-April when late-stage juveniles are present in the population, following each species’ annual birth pulse (**Figure 1B**).

**Figure 1.**
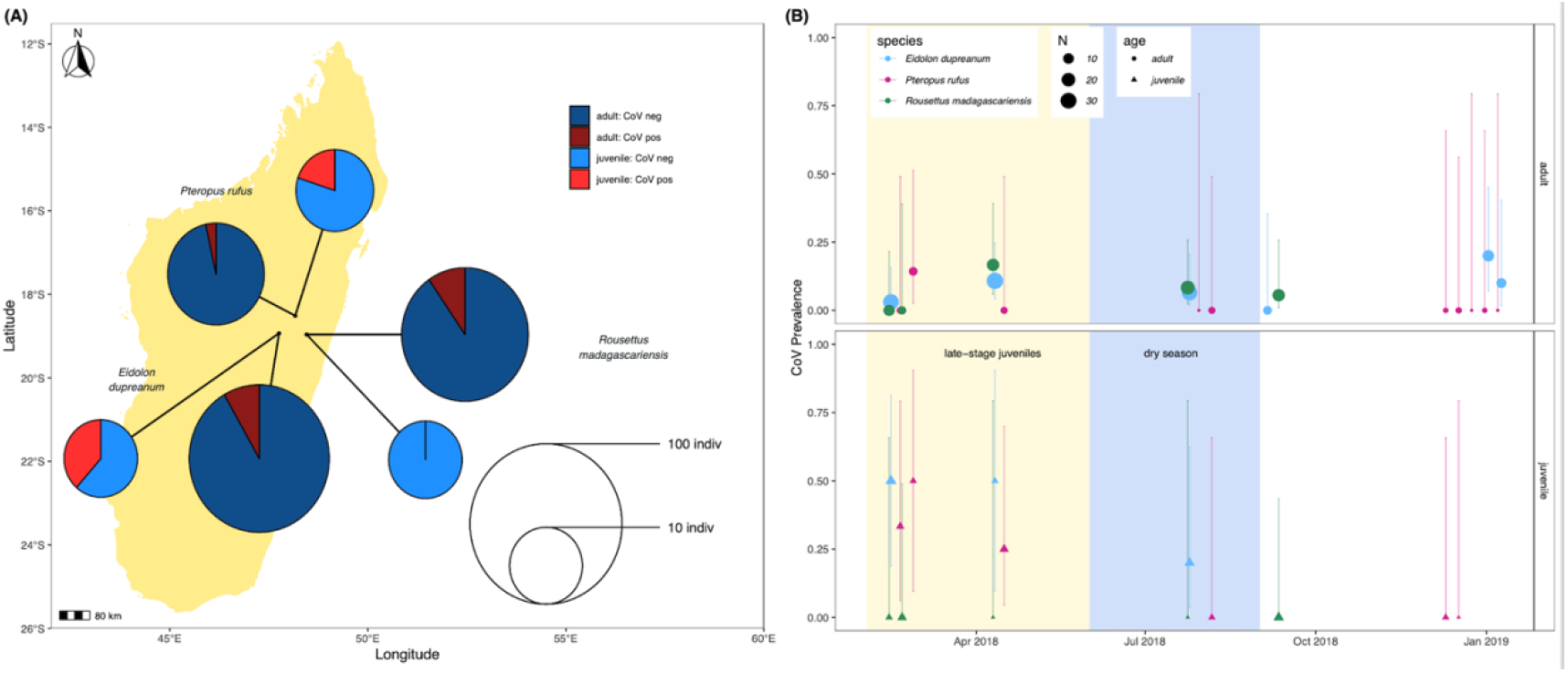
**(A)** Map of sampling sites for *P. rufus, E. dupreanum*, and *R. madagascariensis* in the Districts of Moramanga and Manjakandriana, Madagascar (*P. rufus:* Ambakoana roost; *E. dupreanum*: Angavobe/Angavokely caves; *R. madagascariensis*: Maromizaha cave). Pie charts correspond to coronavirus prevalence in juveniles vs. adults across all three species: 3/15 (20%) vs. 1/29 (3.5%) for *P. rufus*, 5/13 (38.5%) vs. 11/132 (8.3%) for *E. dupreanum*, and 0/13 (0%) vs. 8/83 (9.6%) for *R. madagascariensis*. Pie circle size corresponds to sample size on a log-10 scale. **(B)** Seasonal variation in adult (circle) vs. juvenile (triangle) CoV prevalence by species, from sites depicted in (A). Color corresponds to species and point size to sampling number, as indicated in the legend. Background shading corresponds to the season in which late-stage juveniles are present in the population (yellow) preceding the dry season (lightblue).

### Genome Annotation and BLAST

Three full or near-full CoV genome length contigs were recovered from IDseq for *Nobecoviruses* derived from *R. madagascariensis* (two genomes: 28,980 and 28,926 bps in length) and *P. rufus* (one genome: 29,122 bps in length). In all three genomes, we successfully identified ORF1ab (including RdRp) and structural proteins S (spike), E (envelope), M (matrix), and N (nucleocapsid), in addition to accessory genes NS3, NS7a, and NS7b (**Figure 2A**). In keeping with convention outlined in (61), the accessory genes, NS7a and NS7b, were so named based on nucleotide alignment and amino acid identity to homologous proteins in previously described *Nobecoviruses*.

**Figure 2.**
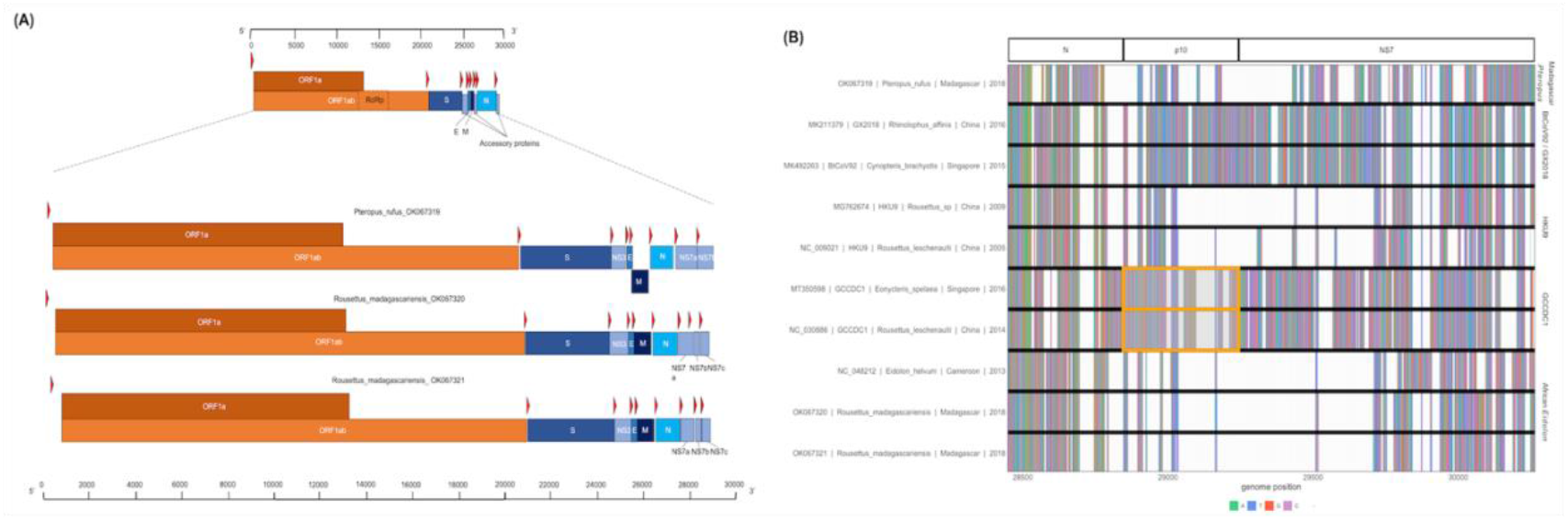
**(A)** Genome structure of novel *Nobecoviruses* derived from *P. rufus* and *R. madagascariensis* fruit bats. TRS locations are highlighted by red arrows, and genes are distinguished by color, with orange corresponding to ORF1a and ORF1b, and various shades of blue to structural proteins S, E, M, and N. Accessory genes NS3, NS7a, NS7b, and NS7c (*R. madagascariensis* genomes only) are depicted in powder blue. **(B)** Multiple sequence alignment of representative sequences from the five main *Nobecovirus* clades, spanning nucleotide positions 28449-30263. This region includes part of the N gene for all sequences and spans the region of p10 orthoreovirus insertion in GCCDC1 lineage (orange highlight), through the NS7 gene region to the 3’ end of each genome.

BLAST analysis of the full genome indicated that the *P. rufus Nobecovirus* sequence is highly divergent, demonstrating only 72.87-73.54% identity to all previously described *Nobecovirus* clades, with the top blast association to *E. helvum Nobecovirus* lineages (**Supplementary Table 1**). Additionally, *Nobecovirus* genomes derived from *R. madagascariensis* demonstrated high identity (∼95%) to *E. helvum Nobecovirus* lineages circulating in Africa. BLASTx analysis of individual genes from viruses derived from both Madagascar species demonstrated the highest identity with previously described *Nobecovirus* sequences in the Orf1ab region (which includes RdRp) for both *P. rufus* and *R. madagascariensis* viruses (76.1% identity for *P. rufus Nobecovirus* to *E. helvum* bat coronavirus and ∼99% identity for *R. madagascariensis Nobecovirus* to *E. helvum* bat coronavirus). By contrast, both *P. rufus* and *R. madagascariensis Nobecovirus* genomes demonstrated substantial divergence from all known homologs in the S gene, where only 46.93-86.9% identity was observed. The *P. rufus Nobecovirus* was similarly divergent in the N gene, though *R. madagascariensis Nobecoviruses* demonstrated high (∼91%) identity to CoV genotypes from *E. helvum* in this region.

In general, BLASTx queries of NS7 accessory proteins in both *R. madagascarienis* and *P. rufus Nobecovirus* demonstrated ∼40-91% amino acid identity to already-characterized *Nobecovirus* proteins (**Supplementary Table 1**). Genomes derived from *R. madagascariensis* appeared slightly more complex than those derived from *P. rufus*, allowing for annotation of one additional accessory gene, NS7c, which has been characterized in recombinant *Nobecovirus* sequences of the RoBat-CoV GCCDC1 lineage (27,61). Curiously, BLASTx query of the NS7a accessory protein in the *P. rufus* genome showed no identity to any previously described *Nobecovirus* protein; rather, the highest scoring protein alignment (31.25% identity, 1e-06 E-value) of the NS7a translation encompassed 40% of the query (query coverage was located at the 3’ end of the query length), and corresponded to an arachnid Low-Density Lipoprotein Receptor-Related Protein 1 (LRP-1) (**Supplementary Table 2**). As LRP-1 is involved in the mammalian innate immune response(88), we hypothesized that this putative novel ORF could be a viral gene involved in immune antagonism. To check the integrity of our *de novo* assembly in NS7a, we mapped the deduplicated raw reads from mNGS to the full genome *P. rufus Nobecovirus* contig generated by IDseq (74) (**Supplementary Figure 1**). We confirmed >200x read coverage across the region corresponding to the putative NS7a accessory protein, with good representation of both forward and reverse-facing reads across the length of the protein, as well as the intergenic regions preceding and succeeding it. We were also able to identify a putative Transcription Regulatory Sequence (TRS) preceding this gene (**Table 1**), further validating our confidence that *P. rufus Nobecovirus* NS7a represents a real though highly divergent protein.

**Table 1.**
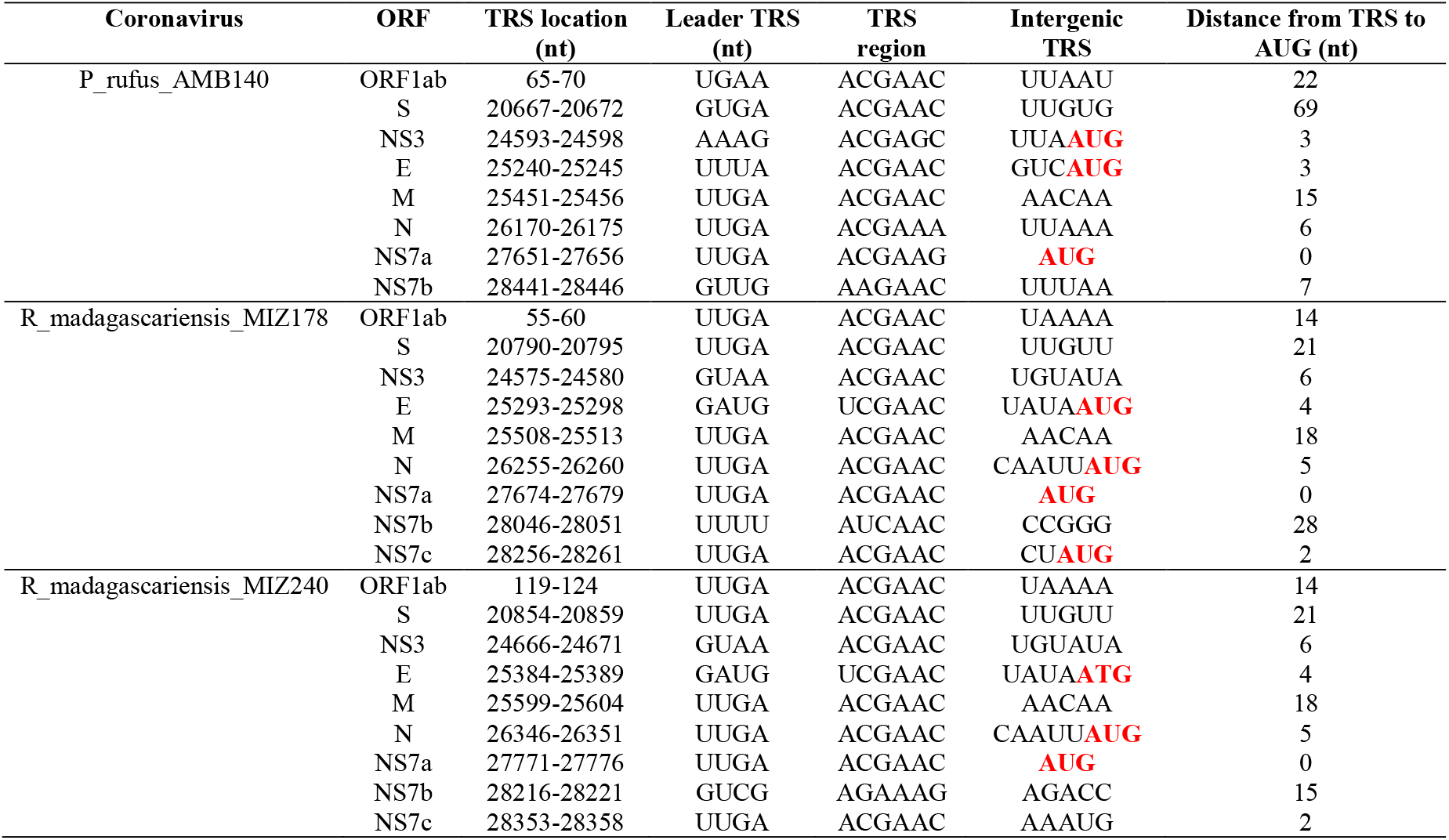
Putative Transcription Regulatory Sequences in novel *Nobecoviruses* from Madagascar fruit bats.

The p10 orthoreovirus insertion within the RoBat-CoV GCCDC1 *Nobecovirus* lineage was not observed in either *Nobecovirus* genomes from *R. madagascariensis* or *P. rufus*. Nonetheless, examination of the multiple sequence alignment of representative sequences of all *Nobecovirus* clades in this region demonstrated the presence of some variable genetic material downstream from the N gene and upstream from the NS7a gene in the divergent *P. rufus Nobecovirus* genome (**Figure 2B**). *Nobecoviruses* clustering in the BtRt-BetaCoV/GX2018 - BtCoV92 lineage also carry a unique coding sequence in this region, highlighting the dynamic nature of the 3’ end of the CoV genome (59).

In addition to the identification of both canonical and novel ORFs described above, we also observed non-coding TRS elements preceding all the major proteins in all three *Nobecovirus* genomes (**Table 1**). Many of these correspond to the 5’-ACGAAC-3’ six bp core motif common to many *Betacoronaviruses*, including SARS-CoV and previously described in *Nobecoviruses* of the GCCDC1 and GX2018/BtCoV92 lineages (58,61,89). For most genes, these TRS elements were located a short distance upstream from the corresponding gene (**Table 1**). Elements identified in the two *R. madagascariensis* genomes were largely comparable, suggesting that these two sequences could represent slight variations in the same virus lineage. Some putative TRS elements, including that preceding *P. rufus* NS7a, showed variation from the 5’-ACGAAC-3’ core motif, with some recapitulating the 5’-AAGAA-3’ motif common to SARS-CoV-2 (90). TRS variations may be indicative of variation in gene expression across individual bats and/or species.

### Phylogenetic Analysis

Phylogenetic analysis of full length *Betacoronavirus* genomes confirmed that both *P. rufus* and *R. madagascariensis* genomes cluster in the *Nobecovirus* subgenus of the *Betacoronaviruses*, with the divergent *P. rufus* forming its own distinct clade and both *R. madagascariensis* genomes grouping with the previously described *E. helvum* reference sequence from Cameroon (60) (**Figure 3A**). We observed distinct groupings of five main *Nobecovirus* lineages in our phylogeny: (a) the largely Asian-derived HKU9 sequences, (b) the African *E. helvum-*derived sequences (now including new *R. madagascariensis Nobecovirus* genomes), (c) the recombinant GCCDC1 genomes, (d) the BtRt-BetaCoV/GX2018 and BtCoV92 genomes described respectively from China and Singapore, and (e) the divergent *P. rufus* genome contributed here from Madagascar. Intriguingly, the *P. rufus* genome groups ancestral to all other *Nobecoviruses*, followed by the *E. helvum/R. madagascariensis* African lineage, with the Asian genotypes forming three distinct (and more recent) clades corresponding to genotypes HKU9, GCCDC1, and GX2018 – BtCoV92. Further phylogenetic analysis of a 259bp fragment of the RdRp gene reconfirmed these groupings and suggested the presence of at least two distinct genetic variants within the *P. rufus* lineage (**Figure 3B**). One RdRp fragment derived from feces of the third Malagasy fruit bat, *E. dupreanum*, grouped within the *E. helvum* – *R. madagascariensis* African *Nobecovirus* lineage, consistent with previous reporting (12). Characterization of the full length genome of this virus will be needed to clarify whether it represents a genetic variant of or a distinct genotype from the *R. madagascariensis* virus. Phylogenetic analysis of the RdRp fragment allowed for inclusion of one partial *Nobecovirus* sequence derived from *E. helvum* bats in Kenya (HQ728482), which also grouped within the *E. helvum* – *R. madagascariensis* African clade, confirming the distribution of this genotype across West and East Africa and into the South-Western Indian Ocean Islands. Notably, one partial Cameroonian *E. helvum* sequence (MG693170) clustered with HKU9 sequences from Asia, rather than within the *E. helvum* – *R. madagascariensis* African clade. These findings suggest that both “African” and “Asian” *Nobecovirus* lineages are likely broadly geographically distributed.

**Figure 3.**
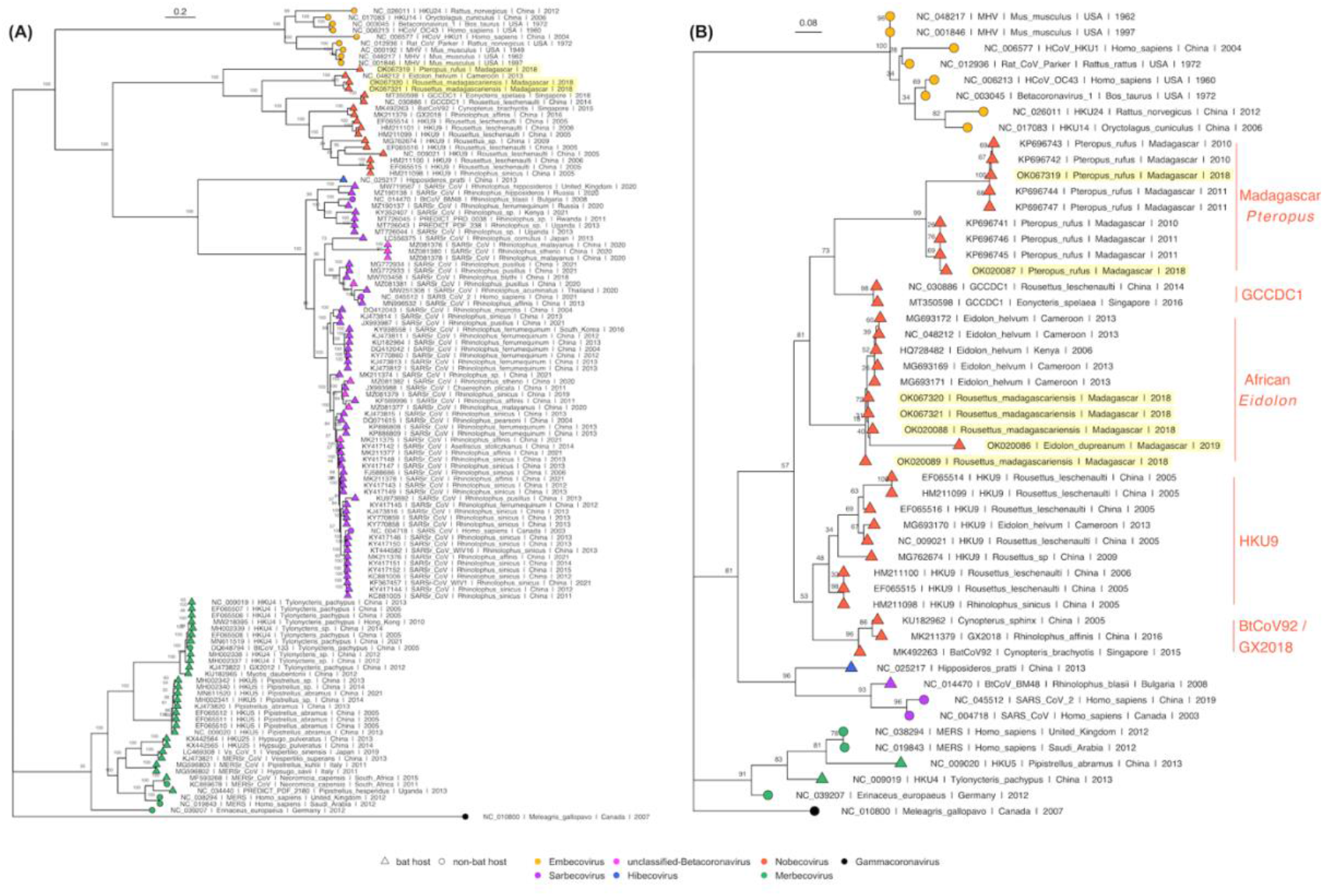
**(A)** Maximum Likelihood phylogeny of full genome *Betacoronavirus* sequences, (RAxML-NG, GTR+I+G4) and **(B)** RdRp phylogeny of a 259bp fragment of *Betacoronavirus* Orf1b (RAxML-NG, TVM+I+G4) (78,79). Bootstrap support values computed using Felsenstein’s method (80) are visualized on tree branches. In both (A) and (B), novel Madagascar sequences are highlighted in yellow, and tip points are colored by *Betacoronavirus* subgenus, corresponding to the legend. Tip shape indicates whether the virus is derived from a bat (triangle) or non-bat (circle) host. Both trees are rooted in turkey *Gammacoronavirus*, accession number NC_010800. Branch lengths are scaled by nucleotide substitutions per site, corresponding to the scalebar given in (A) and (B).

Amino acid phylogenies computed from translated protein alignments of the S, E, M, and N *Betacoronavirus* structural genes (**Supplementary Figure 2**) further confirmed evolutionary relationships suggested in Figure 3. S, M, and N gene phylogenies demonstrated distinct groupings of five main *Nobecovirus* lineages outlined above, while in the E gene phylogeny, the *P. rufus* sequence grouped adjacent to the single Cameroonian-derived *E. helvum* sequence within the HKU9 clade.

Consistent with previous *Betacoronavirus* studies, the UCED molecular clock offered the best fit to the data in our Bayesian analysis of the RdRp fragment. This analysis recapitulated ML support for the five distinct *Nobecovirus* subclades and indicated a time to MRCA for the Madagascar *P. rufus* subclade of ∼1854 (95% HPD: 1505-1999) (**Figure 4**). Data included in this analysis were insufficiently robust to determine reliable estimates of divergence times for the other *Nobecovirus* clades, which showed wide HPD intervals.

**Figure 4.**
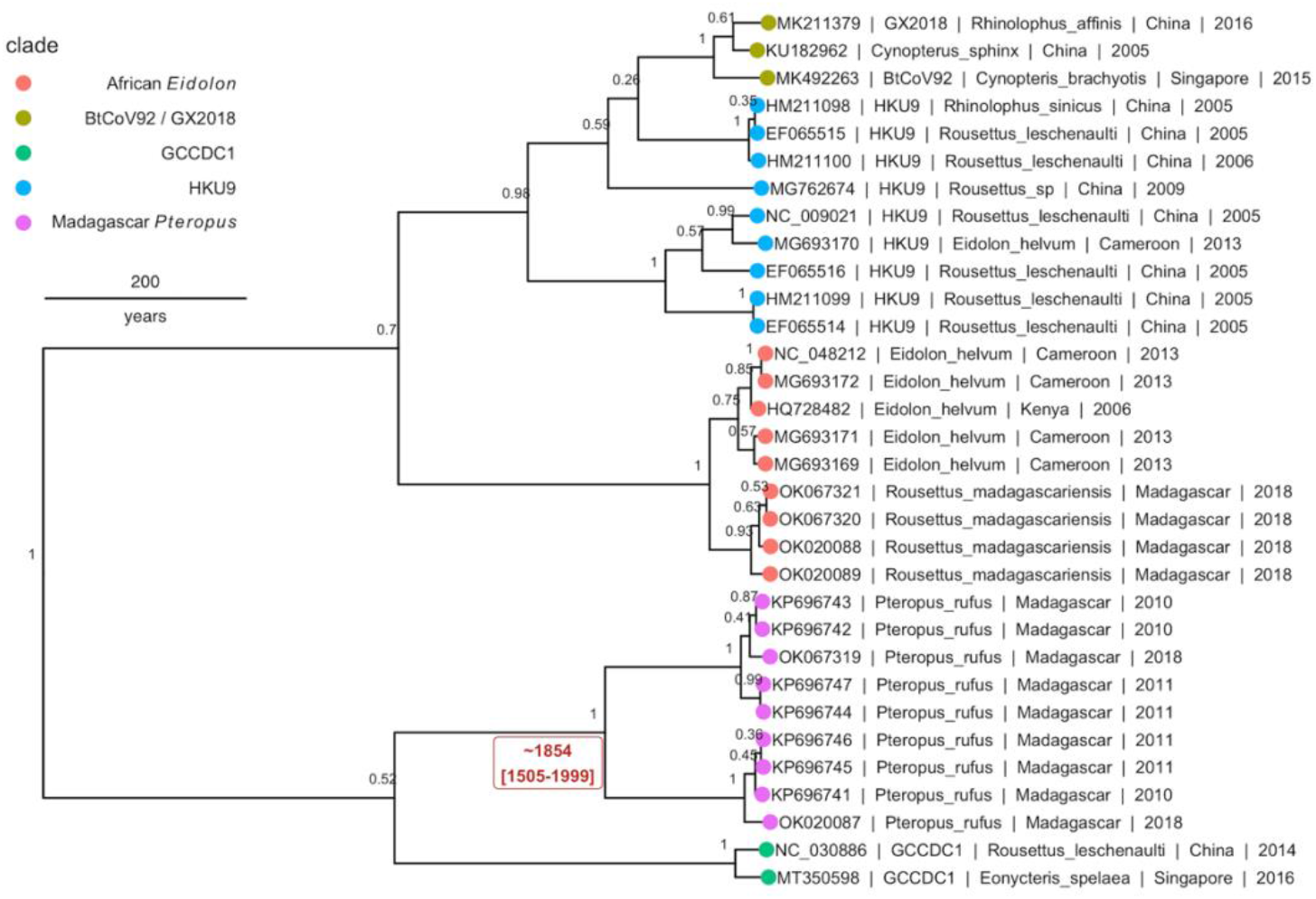
Bayesian phylogeny to estimate time to MRCA for novel *P. rufus Nobecovirus* subclade. Plot depicts output of 1 billion runs of an uncorrelated exponentially distributed relaxed molecular clock Bayesian Skyline Coalescent model (TIM2+G4) as implemented in BEAST2 (82,83). The five major *Nobecovirus* subclades are depicted based on colored tip points, and the mean posterior estimates from averaging of all 1 billion trees after removal of 10% burn-in are visualized above the corresponding node. The date of estimated time to MRCA for the *P. rufus Nobecovirus* subclade (1854) along with the 95% HPD range (1505-1999) is highlighted in red text.

### Recombination Analysis

SimPlot analysis confirmed the evolutionary distinctiveness of the *P. rufus Nobecovirus* genome, which showed <70% amino acid similarity and <50% nucleotide similarity to HKU9, *E. helvum*, and *R. madagascariensis* genotypes across the majority of its genome length (**Figure 5A/B**). Consistent with BLAST results, the *P. rufus Nobecovirus* genome demonstrated the highest similarity to previously described sequences in the Orf1b region, which includes RdRp. The *R. madagascariensis Nobecoviruses*, by contrast, showed >90% amino acid and nucleotide similarity to the *E. helvum* African lineage throughout Orf1ab, but both *P. rufus* and *R. madagascariensis* sequences diverged from all other reference genomes in the first half of the spike protein, which corresponds to the S1 subunit and includes the receptor binding domain that mediates viral entry into host cells (91). Further divergence for both *P. rufus* and *R. madagascariensis Nobecoviruses* was observed in the N structural protein and in the NS7 accessory genes. Bootscan analysis further confirmed these findings, showing that the *P. rufus Nobecovirus* clusters with HKU9 lineages across Orf1ab, NS3, E, and M genes but demonstrates evidence of recombination with *E. helvum* – *R. madagascariensis* African lineages in the S (particularly S1), N, and NS7 genes (**Figure 5C**). Similarly, bootscanning demonstrated that *R. madagascariensis Nobecoviruses* group with the *E. helvum* lineage across Orf1ab, NS3, E, and M but show evidence of recombination with HKU9 and *P. rufus Nobecovirus* in S (again, particularly S1), N, and NS7 genes (**Figure 5C**), thus highlighting the dynamic nature of these regions of the *Nobecovirus* genome.

**Figure 5.**
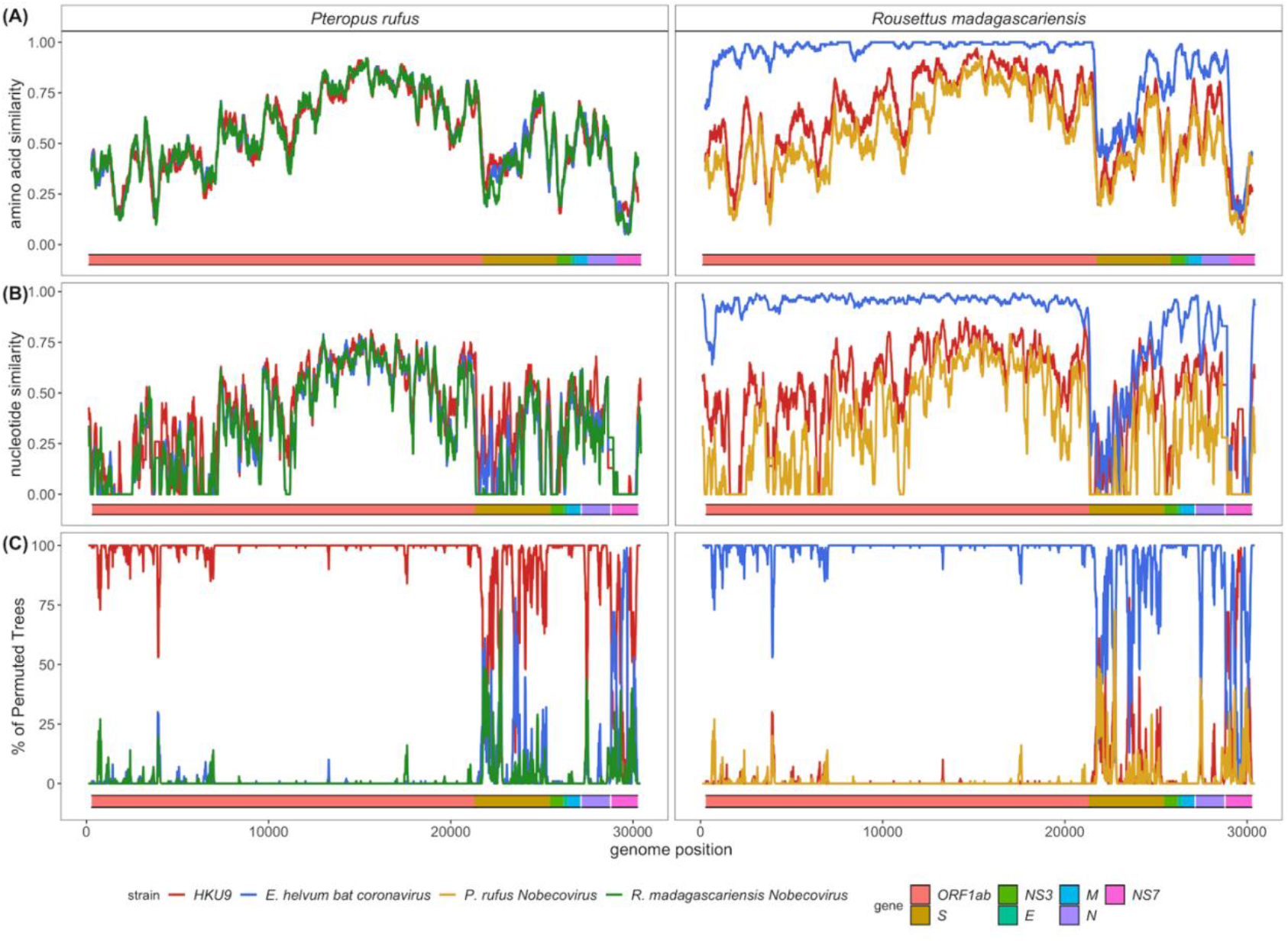
**(A)** Amino acid similarity, **(B)** nucleotide similarity and **(C)** Bootscan plots computed in pySimplot (87) (A) and SimPlot (v.3.5.1) (B and C), using a query sequence of *P. rufus* (left) and *R. madagascariensis* (right)*-* derived *Nobecovirus* sequences. (A) Amino acid similarity plots compares *P. rufus Nobecovirus* and *R. madagascariensis* MIZ240 against one HKU9 (NC_009021) and one *E. helvum* bat CoV (NC_048212) sequence and against each other. Nucelotide similarity and bootscan plots compare *P. rufus Nobecovirus* and both *R. madagascariensis Nobecovirus* sequences against grouped reference sequences corresponding to HKU9 (EF065514-EF065516, HM211098-HM211100, MG693170, NC_009021, MG762674) and *Eidolon helvum* Africa-derived (MG693169, MG693171-MG693172, NC_048212) *Nobecovirus* lineages. Line color indicates similarity (A and B) and bootscan grouping (C) of the query sequence with the corresponding *Nobecovirus* genotype, along disparate regions of the CoV genome, as indicated by the colored bar at the bottom of each plot. Amino acid similarity plots (A) were generated using a window size of 100aa and a step size of 20aa. Nucleotide similarity and bootscan plots (B and C) were generated using a window size of 200bp and a step size of 20bp.

## Discussion

Here, we contribute three full-length genome sequences and four RdRp fragments to public NCBI repositories; these sequences correspond to at least two novel *Nobecoviruses* derived from wild Malagasy fruit bats, *Pteropus rufus* and *Rousettus madagascariensis*, with evidence of additional genetic variants circulating in *Eidolon dupreanum*, as well. Phylogenetic analyses suggest that previously described *Nobecoviruses* can be grouped into five general clades: (a) the HKU9 lineage of largely Asian origins, (b) the mostly African-distributed lineage derived from *E. helvum* bats (which contains our *R. madagascariensis* and *E. dupreanum* sequence contributions), (c) the recombinant GCCDC1 lineage, which has been previously reported from China and Singapore (27,61), (d) the BtRt-BetaCoV/GX2018 – BtCoV92 lineage, also known from China and Singapore (58,59), and (e) a novel, divergent clade corresponding to the newly-described *P. rufus* genome. This new *P. rufus Nobecovirus* has an estimated divergence time from the other *Nobecovirus* subclades within the last 500 years, well after the estimated late Pleistocene divergence of the *P. rufus* host from other Southwest Indian Ocean Island pteropodids (55). Importantly, though largely characterized in Asia, HKU9 *Nobecovirus* genotypes have been identified in West Africa (Cameroon) (60), and *E. helvum* lineages have been characterized across both West (Cameroon) (60) and East (Kenya) (29,92) Africa, as well as on one Indian Ocean island (Madagascar). These findings suggest that different *Nobecovirus* clades may be more broadly geographically distributed than has been previously recognized. To our knowledge, no *Nobecoviruses* have been identified from the southern extension of the pteropodid fruit bat range in Australia; characterization of any CoVs infecting these bats, which are known to host important, zoonotic henipaviruses (93) and lyssaviruses (93), would do much to enhance our understanding of the phylogeography of the *Nobecovirus* clade. Madagascar represents a unique phylogeographic melting pot, with flora and fauna—and corresponding viruses—of both African and Asian descent (51), offering opportunities for mixing of largely disparate viral groups. This mixing is important in light of the CoV penchant for recombination, which can allow viruses from one clade to gain function through acquisition of genetic material from another, thus facilitating rapid changes in host range (39–44).

*Nobecoviruses* are not known to be zoonotic and have been thus far described exclusively infecting fruit bats hosts of the Old World bat family, Pteropodidae. Nonetheless, the *Nobecovirus* subgenus demonstrates a CoV-characteristic tendency to recombine, as evidenced by circulation of the widespread GCCDC1 lineage in Asia, which carries a p10 gene insertion derived from an orthoreovirus between the N structural protein and NS7a accessory protein towards the 3’ end of the genome (61). This orthoreovirus insertion within the GCCDC1 virus genome was not detected among the CoVs in our dataset, though, anecdotally, mNGS of fecal, throat, and urine samples collected in our sampling did identify evidence of orthoreovirus infection in several throat swabs derived from *E. dupreanum* bats, highlighting the potential for recombination opportunities between these two viral groups. Notably, recombination analyses suggested substantial selection has taken place in this region of both *R. madagascariensis* and *P. rufus-*derived *Nobecoviruses*. Selection at the 3’ end of the CoV genome may modulate viral replication ability, since several regulatory sequences and accessory genes (e.g. NS7) are defined in this region (94). Viral replication ability may be further impacted by variation in TRS motifs, which regulate expression of corresponding genes. We identified putative TRS sequences corresponding to all structural and non-structural genes identified in all three contributed *Nobecovirus* genomes; while the majority of these TRS motifs recapitulated the well-conserved 5’-ACGAAC-3’ *Betacoronavirus* core sequence (58,61), variation in a subset of genes across species and individuals (e.g. differing motifs between two *R. madagascariensis*-derived genomes) may correspond to variation in gene expression.

Recombination potential is a particular cause for concern in cases where viruses that lack the ability to infect human cells may acquire this zoonotic capacity through genetic exchange with other viruses coinfecting the same host. Indeed, the original SARS-CoV is believed to have acquired its capacity to bind human ACE2 through a recombination event with ACE2-using *Sarbecoviruses* in the disparate SARS-CoV-2 clade (94). *Sarbecoviruses*, in particular, are known to recombine frequently, giving rise to new genetic variants, in regions where different species of Rhinolophid bat hosts co-roost and share viruses (7). Cave-resident Malagasy fruit bats, *E. dupreanum* and *R. madagascariensis*, are known to co-roost with each other and with several species of insectivorous bat (63), which could facilitate *Nobecovirus* recombination. The observed similarity in *Nobecovirus* sequences derived from *E. dupreanum* and *R. madagascariensis* (which cluster in the same lineage), as compared with disparate sequences derived from tree-roosting *P. rufus*, suggest that some CoV genetic exchange may have already taken place between bats with overlapping habitats. To date, zoonotic potential has not been demonstrated for any previously described *Nobecoviruses*, and Rhinolophid bats associated with ACE2 usage are not resident in Madagascar. Nonetheless, bats in family Vespertilionidae, the family most commonly associated with zoonotic *Merbecoviruses* (8–10), are widespread in Madagascar, and *Mormopterus jugularis*, a known Molossidae bat host for *Alphacoronaviruses* of undetermined zoonotic potential (32), has been described co-roosting with *R. madagascariensis* (95). Bootscan analyses identified signatures of recombination in the S1 subunit of both *P. rufus* and *R. madagascariensis Nobecovirus* spike proteins, suggesting that this region of the genome, which modulates host range through cell surface receptor binding, may be under selective pressure.

In addition to posing risk for future zoonoses, *Nobecoviruses* derived from wild Madagascar fruit bats could provide unprecedented genetic material for recombination to existing human coronaviruses already in circulation across the island—most notably SARS-CoV-2 (71). At the time of this writing, COVID-19 infections remain widespread and vaccination limited across Madagascar (96). Previous work has assessed the risk of reverse zoonosis, or ‘spillback’ of SARS-CoV-2 from human to bat populations in the United States (16), concluding that high human caseloads and frequent human-bat contact rates in research settings pose both conservation risks to naïve bat populations presented with a novel pathogen, as well as human health risks presented by the possible establishment of secondary wildlife reservoirs for SARS-CoV-2 capable of sourcing future epidemics or the generation of unique viral variants through human-wildlife virus recombination (16). Bat-human contact rates are higher, on average, in Madagascar than in the US, as bats are consumed across the island for subsistence and frequently found roosting in human establishments or human-adjacent habitats (64–68). SARS-CoV-2 has already demonstrated its capacity for successful reverse zoonosis and adaptation to non-human hosts, in the case of farmer-sourced infections of mink in Finland (97), underscoring the legitimacy of these concerns. Notably, spillback is less likely to be an issue in regions where animals are killed upon capture for consumption (vs. transported live), as is often the case in Madagascar (68).

Prevalence of coronavirus RNA by sequence detection in fecal samples averaged around 10% across all three Malagasy fruit bat species examined in our study, consistent with CoV prevalence reported in wild bat species elsewhere (12,32). One previous study of CoV circulation in Madagascar fruit bats reported much lower prevalence of infection in *E. dupreanum* and *R. madagascariensis*-derived fecal specimens, respectively 1/88 (1.1%) and 0/141 (0%), as compared with a 13/88 (14.8%) prevalence in *P. rufus-*derived feces (12). As in our study, this previous work found no positive infections in throat swabs, supporting a gastrointestinal tropism for CoVs in this fruit bat system, in contrast to the respiratory infections more commonly observed in humans. One additional study in the West Indian Ocean provided more information about CoV prevalence in Madagascar bats, with 6/45 (13.3%) *R. madagascariensis* fecal specimens testing CoV positive, as compared to 10/63 (15.9%) *M. jugularis* specimens, 4/44 (9.1%) *Triaenops menamena* specimens, and 2/21 (9.5%) *Mops midas* specimens (32). Consistent with previous findings (31,38,98,99), we observed the highest prevalence of CoV infection in *P. rufus* and *E. dupreanum* juveniles. We hypothesize that the absence of juvenile infection identified in *R. madagascariensis* bats in our study could be due to the staggered nature of the birth pulse for these three species: Madagascar fruit bats birth in three successive birth pulse waves, led by *P. rufus* in October, and followed by *E. dupreanum* in November and *R. madagascariensis* in December and January (100). As the bulk of juvenile *R. madagascariensis* bats sampled our study were captured in February, it is possible that most were still too young to be CoV-positive (perhaps under protection from inherited maternal immunity (57)). By the time of the second *R. madagascariensis* sampling in April, juveniles would have been large enough to be erroneously classed as adults, as size range variation is more limited in small *R. madagascariensis* bats as compared with the two other Malagasy fruit bat species (63).

Our work emphasizes the importance of longitudinal ecological studies in identifying viral shedding events in transiently-infected wildlife hosts across multiple age and reproductive classes. Enhanced future surveillance efforts will be useful in pinpointing the exact seasonality of peak CoV shedding events, and mitigation efforts for both zoonotic and reverse zoonotic risks should be focused on limiting human-bat contact (in particular, the government-sanctioned hunting seasons) during these periods. Our study highlights the enhanced evolutionary and functional virological inference that can be derived from full genome sequences, detected by unbiased metagenomic sequencing. Characterization of these genomes provides the basis for basic virology experiments to follow, such as pseudovirus or reverse genetics experiments aimed at understanding host receptor utilization. More thorough studies documenting the seasonal dynamics of bat-borne CoVs, which elucidate genetic variation within and between species that share habitats in wild populations will be essential to understanding CoV recombination, host shifting, and zoonotic potential. Replication of such studies across the global range of both coronaviruses and their bat hosts, in particular in understudied regions of Africa, is needed to assess the landscape of future zoonotic risks and present opportunities for intervention and mitigation.

## Author Contributions

CEB conceived of the project and acquired the funding, in collaboration with JMH, PD, JLD, and CMT. Field samples were collected and RNA extracted by AA, SA, AG, HCR, THR, NAFR, and CEB. AK led the mNGS, with support from VA, HCR, THR, and CEB. GK and CEB analyzed the resulting data and co-wrote the original draft of the manuscript, which all authors edited and approved.

## Acknowledgements

The authors acknowledge Kimberly Rivera and Sarah Guth for help in the field and the lab and the Virology Unit at the Institut Pasteur de Madagascar and Maira Phelps of the Chan Zuckerberg Biohub (CZB) for logistical support. They additionally thank Angela Detweiler, Michelle Tan, and Norma Neff of the CZB genomics platform for mNGS support and thank the Brook lab at the University of Chicago for helpful contributions to the manuscript.

## Data Availability Statement

All sequencing data has been deposited for public access on NVBI Virus (accession numbers OK020086-OK020089 and OK067319-OK067321). All code and metadata used in all analyses is available for download from our public Github repository at: https://github.com/brooklabteam/Mada-Bat-CoV/.

## Supplementary Materials

**Supplementary Figure 1.**
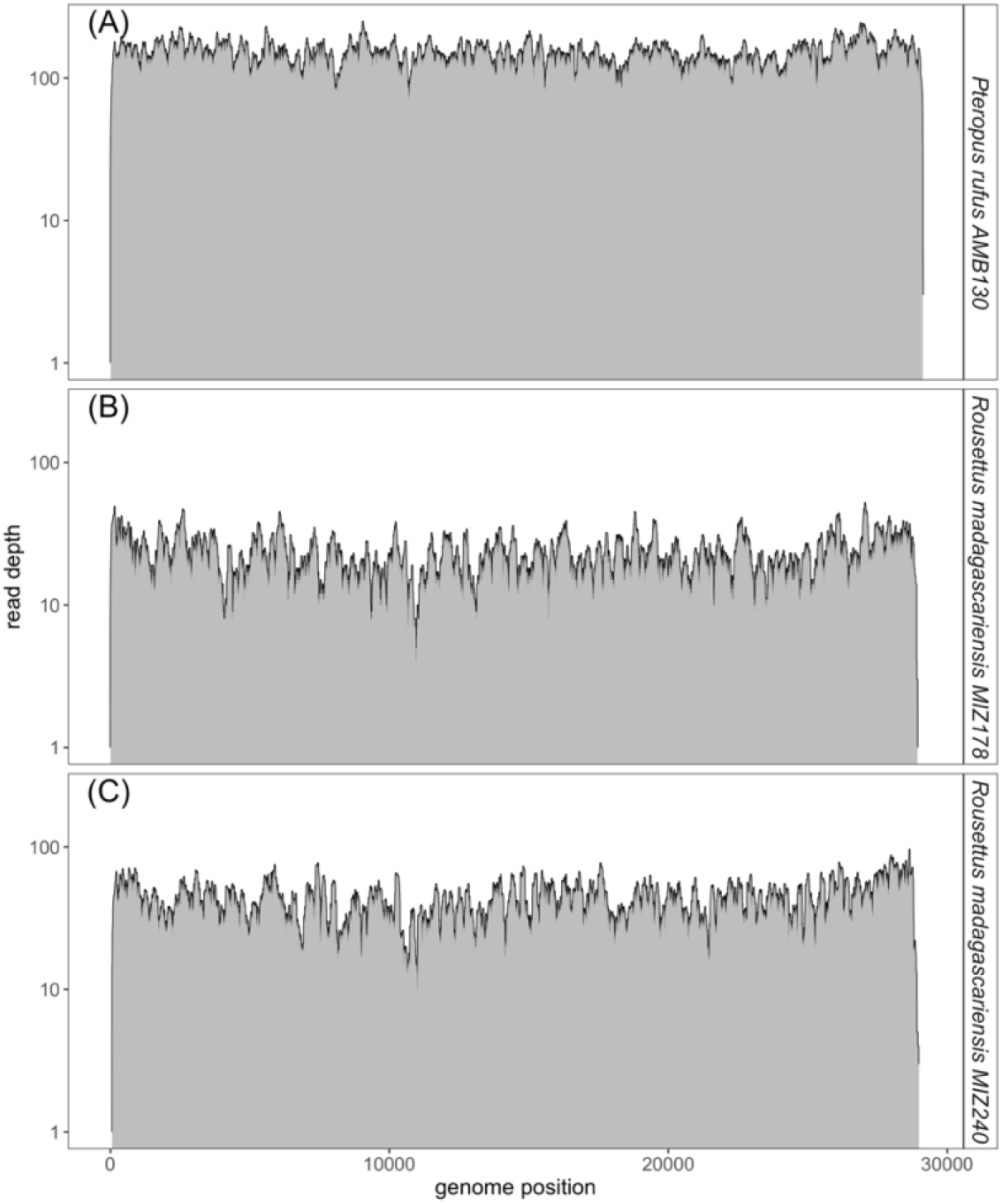
Read depth and coverage for full genome *Nobecovirus* sequences identified in Malagasy fruit bats. Read depth after deduplication by CDHIT (101) for full genome Madagascar fruit bat *Nobecoviru*s contigs assembled in IDseq (74). Viral genomes were assembled from fecal specimens derived from one *P. rufus* (A) and two *R. madagascariensis* (B and C) bats.

**Supplementary Figure 2.**
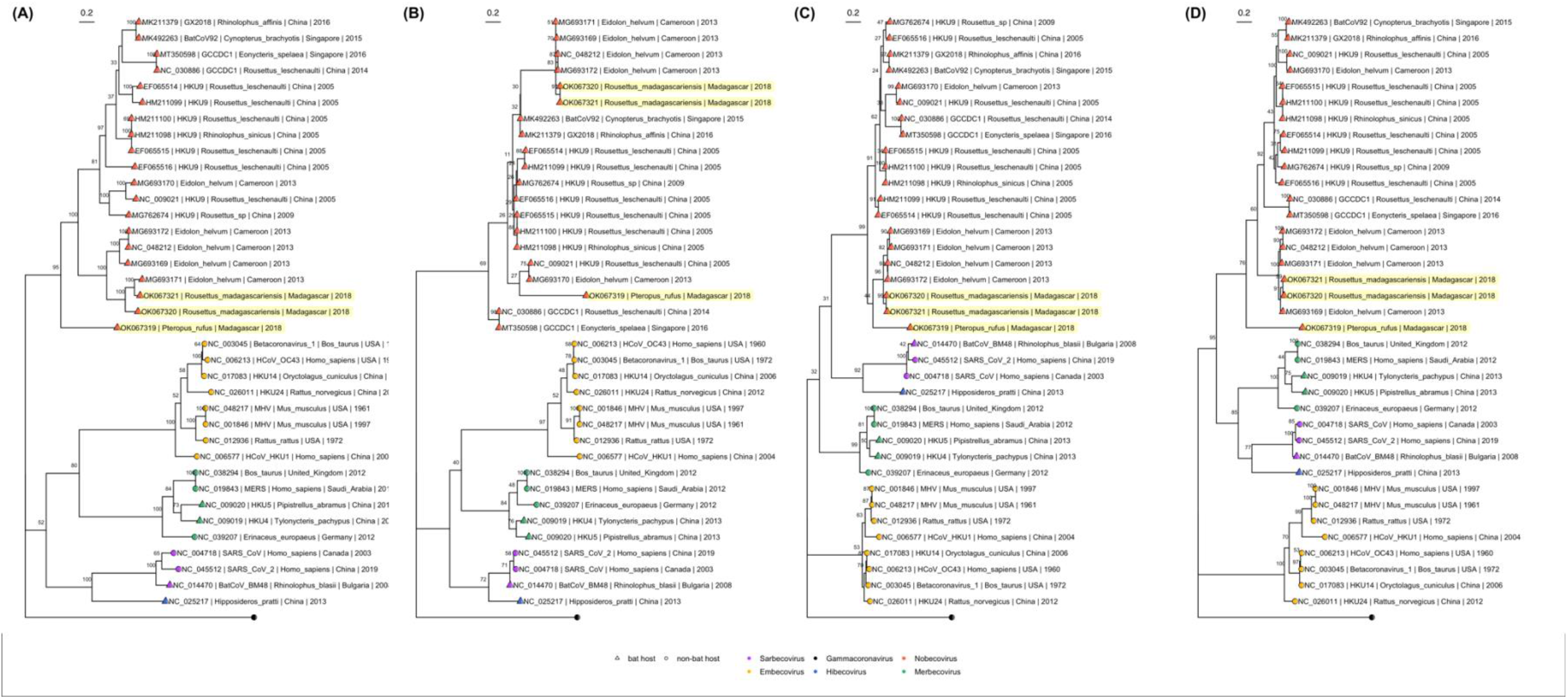
ML amino acid Betacoronavirus phylogenies. Maximum Likelihood amino acid phylogenies corresponding to translated sequences of the **(A)** spike, **(B)** envelope, **(C)** matrix, and **(D)** nucleocapsid *Betacoronavirus* proteins. All phylogenies were computed in RAxML-NG, using respective amino acid substitution models (A) WAG+I+G4+F, (B) LG+G4, (C) LG+I+G4, and (D) LG+I+G4+F (79,80). Bootstrap support values computed using Felsenstein’s method (81) are visualized on tree branches. In (A-D) novel Madagascar sequences are highlighted in yellow, and tip points are colored by *Betacoronavirus* subgenus, corresponding to the legend. Tip shape indicates whether the virus is derived from a bat (triangle) or non-bat (circle) host. Both trees are rooted in turkey *Gammacoronavirus*, accession number NC_010800. Branch lengths are scaled by amino acid substitutions per site, corresponding to the scale bar given indicated in each subplot.

**Supplementary Table 1.**
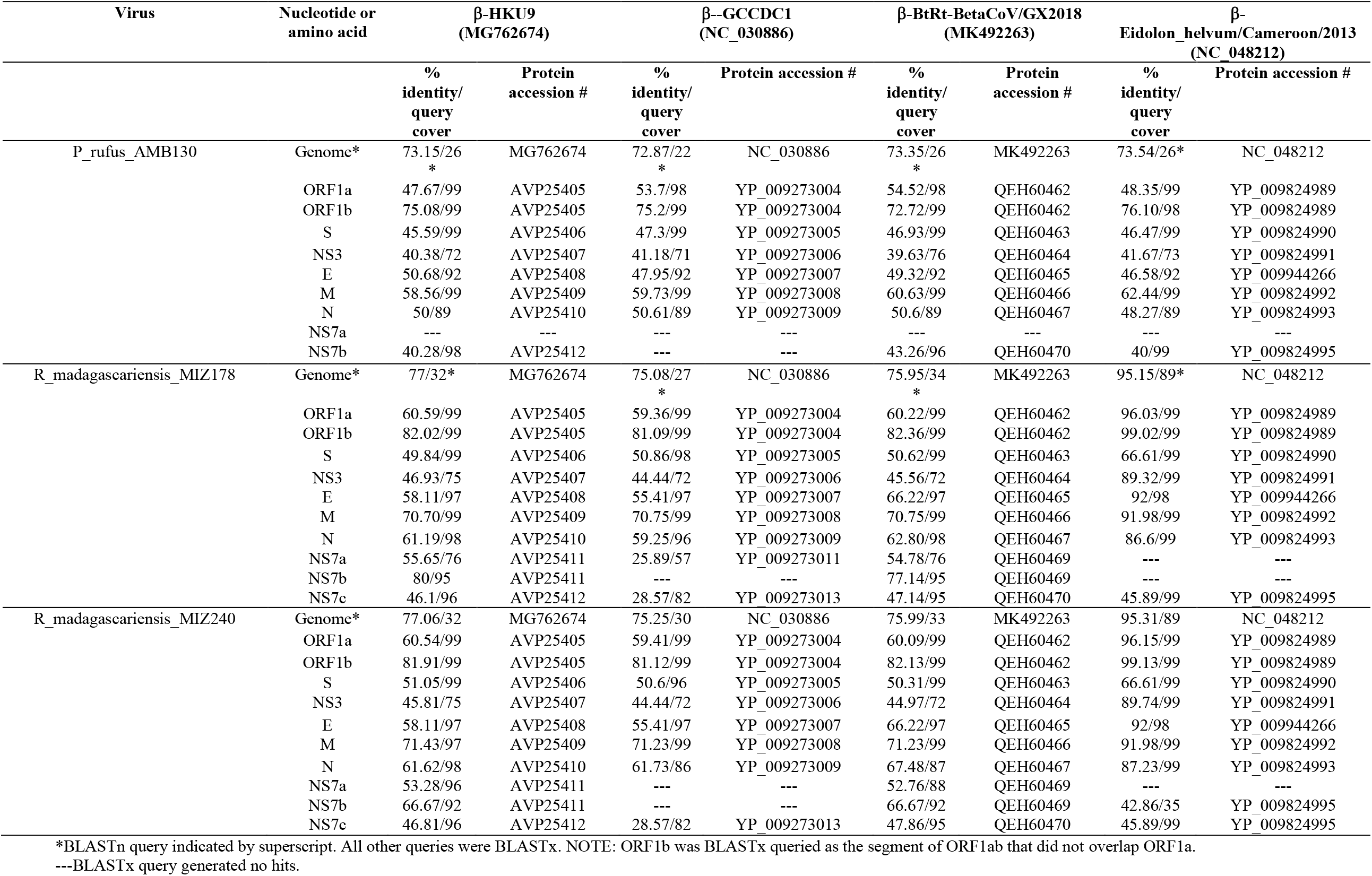
Summary of BLAST queries to reference homologs for proteins identified in Malagasy fruit bat *Nobecoviruses*.

**Supplementary Table 2.**
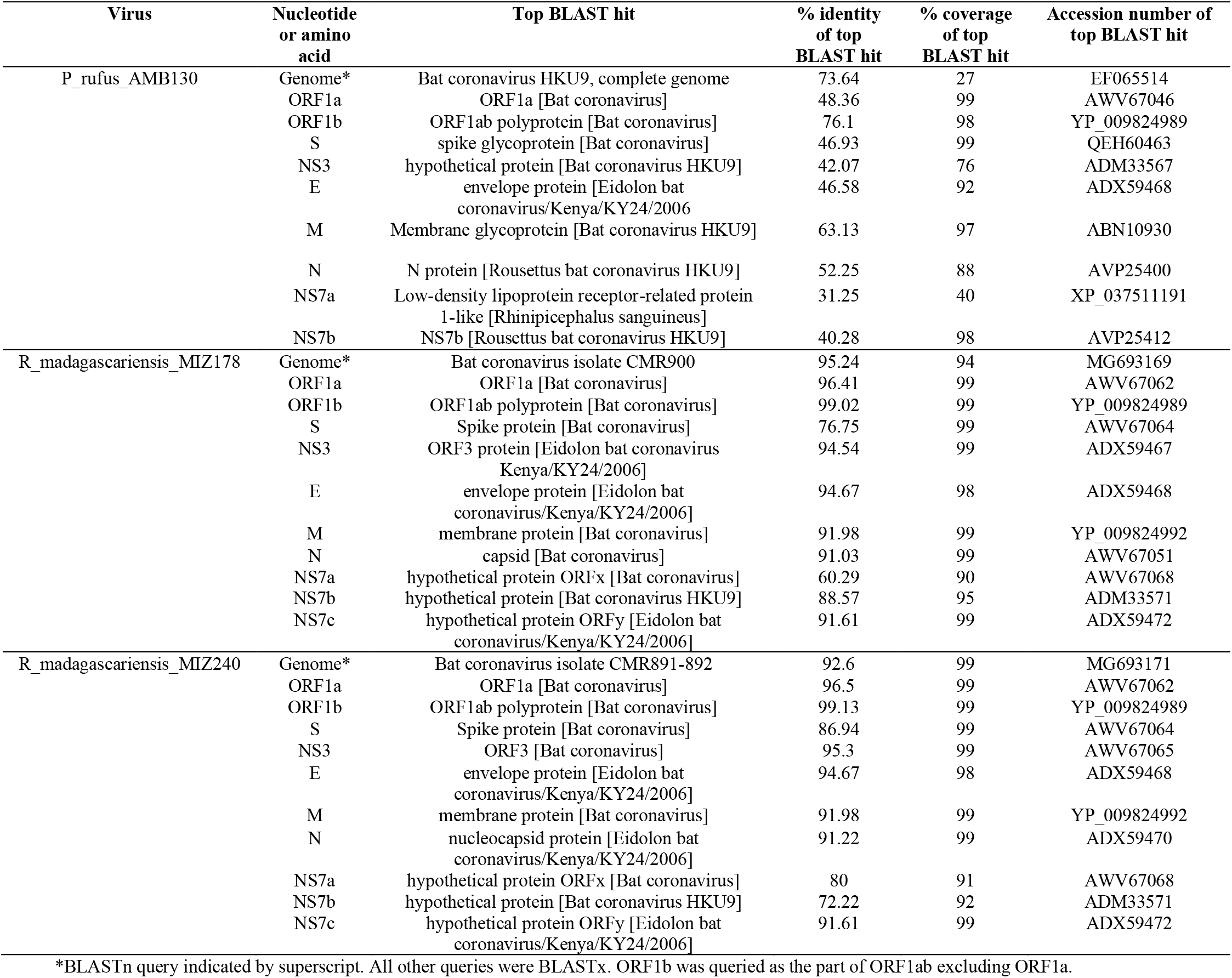
Summary of BLAST queries to reference homologs for proteins identified in Malagasy fruit bat *Nobecoviruses*.

## References

1. Banerjee A, Kulcsar K, Misra V, Frieman M, Mossman K. Bats and coronaviruses. Viruses. 2019;11(1):7–9.

2. Wu F, Zhao S, Yu B, Chen Y-M, Wang W, Song Z-G, et al. A new coronavirus associated with human respiratory disease in China. Nature. 2020;

3. Hu B, Ge X, Wang LF, Shi Z. Bat origin of human coronaviruses. Virology Journal. 2015;12(1):1–10.

4. Ravelomanantsoa NAF, Guth S, Andrianiaina A, Andry S, Gentles A, Ranaivoson HC, et al. The zoonotic potential of bat-borne coronaviruses. Emerging Topics in Life Sciences. 2020 Dec 11;4(4).

5. Wille M, Holmes EC. Wild birds as reservoirs for diverse and abundant gamma- And deltacoronaviruses. FEMS Microbiology Reviews. 2020;44(5):631–44.

6. Drexler JF, Gloza-Rausch F, Glende J, Corman VM, Muth D, Goettsche M, et al. Genomic characterization of severe acute respiratory syndrome-related coronavirus in European bats and classification of coronaviruses based on partial RNA-dependent RNA polymerase gene sequences. Journal of virology. 2010/08/04. 2010 Nov;84(21):11336–49.

7. Hu B, Zeng L-P, Yang X-L, Ge X-Y, Zhang W, Li B, et al. Discovery of a rich gene pool of bat SARS-related coronaviruses provides new insights into the origin of SARS coronavirus. PLoS pathogens [Internet]. 2017 Nov 30;13(11):e1006698–e1006698. Available from: https://pubmed.ncbi.nlm.nih.gov/29190287

8. Anthony SJ, Gilardi K, Menachery VD, Goldstein T, Ssebide B, Mbabazi R, et al. Further evidence for bats as the evolutionary source of middle east respiratory syndrome coronavirus. mBio. 2017;8(2):1–13.

9. Woo PC, Lau SK, Li KS, Tsang AK, Yuen K-Y. Genetic relatedness of the novel human group C betacoronavirus to Tylonycteris bat coronavirus HKU4 and Pipistrellus bat coronavirus HKU5. Emerging microbes & infections. 2012/11/07. 2012 Nov;1(11):e35– e35.

10. Corman VM, Ithete NL, Richards LR, Schoeman MC, Preiser W, Drosten C, et al. Rooting the phylogenetic tree of middle East respiratory syndrome coronavirus by characterization of a conspecific virus from an African bat. Journal of virology. 2014/07/16. 2014 Oct;88(19):11297–303.

11. Frutos R, Serra-Cobo J, Pinault L, Lopez Roig M, Devaux CA. Emergence of bat-related Betacoronaviruses: Hazard and risks. Frontiers in Microbiology. 2021;12:437.

12. Razanajatovo NH, Nomenjanahary LA, Wilkinson DA, Razafimanahaka JH, Goodman SM, Jenkins RK, et al. Detection of new genetic variants of Betacoronaviruses in endemic frugivorous bats of Madagascar. Virology Journal [Internet]. 2015;12(1):42. Available from: https://doi.org/10.1186/s12985-015-0271-y

13. Woo PCY, Huang Y, Lau SKP, Yuen K-Y. Coronavirus genomics and bioinformatics analysis. Viruses [Internet]. 2010/08/24. 2010 Aug;2(8):1804–20. Available from: https://pubmed.ncbi.nlm.nih.gov/21994708

14. Chen S-C, Olsthoorn RCL, Yu C-H. Structural phylogenetic analysis reveals lineage-specific RNA repetitive structural motifs in all coronaviruses and associated variations in SARS-CoV-2. Virus Evolution. 2021 Jan 20;7(1).

15. Zhou H, Chen X, Hu T, Li J, Song H, Liu Y, et al. A novel bat coronavirus reveals natural insertions at the S1/S2 cleavage site of the Spike protein and a possible recombinant origin of HCoV-19. bioRxiv. 2020 Jan 1;2020.03.02.974139.

16. Olival KJ, Cryan PM, Amman BR, Baric RS, Blehert DS, Brook CE, et al. Possibility for reverse zoonotic transmission of SARS-CoV-2 to free-ranging wildlife: A case study of bats. PLOS Pathogens [Internet]. 2020 Sep 3;16(9):e1008758.. Available from: https://doi.org/10.1371/journal.ppat.1008758

17. Forni D, Cagliani R, Sironi M. Recombination and positive selection differentially shaped the diversity of Betacoronavirus subgenera. Viruses. 2020 Nov 16;12(11):1313.

18. Llanes A, Restrepo CM, Caballero Z, Rajeev S, Kennedy MA, Lleonart R. Betacoronavirus genomes: How genomic information has been used to deal with past outbreaks and the COVID-19 pandemic. International journal of molecular sciences. 2020 Jun 26;21(12):4546.

19. Li W, Shi Z, Yu M, Ren W, Smith C, Epstein JH, et al. Bats are natural reservoirs of SARS-like coronaviruses. Science. 2005 Oct 28;310(5748):676.

20. Hul V, Delaune D, Karlsson EA, Hassanin A, Tey PO, Baidaliuk A, et al. A novel SARS-CoV-2 related coronavirus in bats from Cambodia. bioRxiv. 2021 Jan 1;2021.01.26.428212.

21. Valitutto MT, Aung O, Tun KYN, Vodzak ME, Zimmerman D, Yu JH, et al. Detection of novel coronaviruses in bats in Myanmar. PLOS ONE. 2020 Apr 9;15(4):e0230802..

22. Lau SKP, Li KSM, Huang Y, Shek C-T, Tse H, Wang M, et al. Ecoepidemiology and complete genome comparison of different strains of severe acute respiratory syndrome-related Rhinolophus bat coronavirus in China reveal bats as a reservoir for acute, self-limiting infection that allows recombination events. Journal of virology. 2010/01/13. 2010 Mar;84(6):2808–19.

23. Latinne A, Hu B, Olival KJ, Zhu G, Zhang L, Li H, et al. Origin and cross-species transmission of bat coronaviruses in China. Nature Communications. 2020;11(1):4235.

24. Wacharapluesadee S, Duengkae P, Rodpan A, Kaewpom T, Maneeorn P, Kanchanasaka B, et al. Diversity of coronavirus in bats from Eastern Thailand. Virology Journal. 2015;12(1):57.

25. Zhou H, Ji J, Chen X, Bi Y, Li J, Wang Q, et al. Identification of novel bat coronaviruses sheds light on the evolutionary origins of SARS-CoV-2 and related viruses. Cell. 2021 Aug;184(17).

26. Lacroix A, Duong V, Hul V, San S, Davun H, Omaliss K, et al. Genetic diversity of coronaviruses in bats in Lao PDR and Cambodia. Infection, Genetics and Evolution. 2017 Mar;48.

27. Paskey AC, Ng JHJ, Rice GK, Chia WN, Philipson CW, Foo RJH, et al. Detection of recombinant Rousettus bat coronavirus GCCDC1 in lesser dawn bats (Eonycteris spelaea) in Singapore. Viruses [Internet]. 2020 May 14;12(5):539. Available from: https://www.ncbi.nlm.nih.gov/pmc/articles/PMC7291116/

28. Anthony SJ, Ojeda-Flores R, Rico-Chávez O, Navarrete-Macias I, Zambrana-Torrelio CM, Rostal MK, et al. Coronaviruses in bats from Mexico. Journal of General Virology. 2013;94(PART 5):1028–38.

29. Tong S, Conrardy C, Ruone S, Kuzmin I v, Guo X, Tao Y, et al. Detection of novel SARS-like and other coronaviruses in bats from Kenya. Emerging infectious diseases [Internet]. 2009 Mar;15(3):482–5. Available from: https://pubmed.ncbi.nlm.nih.gov/19239771

30. Tao Y, Tong S. Complete genome sequence of a Severe Acute Respiratory Syndrome-related Coronavirus from Kenyan bats. Microbiology Resource Announcements. 2019;8(28):e00548–19.

31. Montecino-Latorre D, Goldstein T, Gilardi K, Wolking D, van Wormer E, Kazwala R, et al. Reproduction of East-African bats may guide risk mitigation for coronavirus spillover. One Health Outlook [Internet]. 2020;2(1):2. Available from: https://doi.org/10.1186/s42522-019-0008-8

32. Joffrin L, Goodman SM, Wilkinson DA, Ramasindrazana B, Lagadec E, Gomard Y, et al. Bat coronavirus phylogeography in the Western Indian Ocean. Scientific Reports. 2020;10(1):6873.

33. Su S, Wong G, Shi W, Liu J, Lai ACK, Zhou J, et al. Epidemiology, genetic recombination, and pathogenesis of coronaviruses. Trends in Microbiology. 2016;24(6):490–502.

34. Woolhouse MEJ, Haydon DT, Antia R. Emerging pathogens: the epidemiology and evolution of species jumps. Trends in Ecology and Evolution. 2005 May;20(5):238–44.

35. Zhou P, Yang X-L, Wang X-G, Hu B, Zhang L, Zhang W, et al. A pneumonia outbreak associated with a new coronavirus of probable bat origin. Nature. 2020;

36. Li W, Moore MJ, Vasilieva N, Sui J, Wong SK, Berne MA, et al. Angiotensin-converting enzyme 2 is a functional receptor for the SARS coronavirus. Nature. 2003;147(2):120–1.

37. Raj VS, Mou H, Smits SL, Dekkers DHW, Müller MA, Dijkman R, et al. Dipeptidyl peptidase 4 is a functional receptor for the emerging human coronavirus-EMC. Nature. 2013;495(7440):251–4.

38. Anthony SJ, Johnson CK, Greig DJ, Kramer S, Che X, Wells H, et al. Global patterns in coronavirus diversity. Virus evolution [Internet]. 2017 Jun 12;3(1):vex012–vex012. Available from: https://pubmed.ncbi.nlm.nih.gov/28630747

39. Vijgen L, Keyaerts E, Moes E, Thoelen I, Wollants E, Lemey P, et al. Complete genomic sequence of human coronavirus OC43: Molecular clock analysis suggests a relatively recent zoonotic coronavirus transmission event. Journal of Virology. 2005;79(3):1595– 604.

40. Lau SKP, Lee P, Tsang AKL, Yip CCY, Tse H, Lee RA, et al. Molecular epidemiology of human coronavirus OC43 reveals evolution of different genotypes over time and recent emergence of a novel genotype due to natural recombination. Journal of Virology. 2011 Nov 1;85(21).

41. Woo PCY, Lau SKP, Huang Y, Tsoi H-W, Chan K-H, Yuen K-Y. Phylogenetic and recombination analysis of coronavirus HKU1, a novel coronavirus from patients with pneumonia. Archives of Virology. 2005 Nov 28;150(11).

42. Sabir JSM, Lam TT-Y, Ahmed MMM, Li L, Shen Y, E. M. Abo-Aba S, et al. Co-circulation of three camel coronavirus species and recombination of MERS-CoVs in Saudi Arabia. Science. 2016 Jan 1;351(6268).

43. Wang Y, Liu D, Shi W, Lu R, Wang W, Zhao Y, et al. Origin and possible genetic recombination of the Middle East respiratory syndrome coronavirus from the first imported case in China: Phylogenetics and coalescence analysis. mBio. 2015 Oct 30;6(5).

44. Lau SKP, Feng Y, Chen H, Luk HKH, Yang W-H, Li KSM, et al. Severe Acute Respiratory Syndrome (SARS) coronavirus ORF8 protein is acquired from SARS-related coronavirus from greater horseshoe bats through recombination. Journal of Virology. 2015;89(20):10532–47.

45. Graham RL, Baric RS. Recombination, reservoirs, and the modular spike: mechanisms of coronavirus cross-species transmission. Journal of virology. 2009/11/11. 2010 Apr;84(7):3134–46.

46. Ogando NS, Ferron F, Decroly E, Canard B, Posthuma CC, Snijder EJ. The curious case of the Nidovirus exoribonuclease: Its role in RNA synthesis and replication fidelity. Frontiers in Microbiology. 2019;10:1813.

47. Nga PT, de Parquet MC, Lauber C, Parida M, Nabeshima T, Yu F, et al. Discovery of the first insect nidovirus, a missing evolutionary link in the emergence of the largest RNA virus genomes. PLoS Pathogens. 2011;7(9):e1002215.

48. Gorbalenya AE, Enjuanes L, Ziebuhr J, Snijder EJ. Nidovirales: Evolving the largest RNA virus genome. Virus Research. 2006;117(1):17–37.

49. Smith EC, Blanc H, Vignuzzi M, Denison MR. Coronaviruses lacking exoribonuclease activity are susceptible to lethal mutagenesis: Evidence for proofreading and potential therapeutics. PLoS Pathogens. 2013;9(8).

50. Lai MMC. RNA recombination in animal and plant viruses. Microbiological Reviews. 1992;56(1):61–79.

51. Masters J, Wit M de, Asher RJ. Reconciling the origins of Africa, India and Madagascar with vertebrate dispersal scenarios. Folia Primatologica. 2006;3200:399–418.

52. Species IUCN Red List Threat. IUCN 2018. Version 2018-2.

53. Shi JJ, Chan LM, Peel AJ, Lai R, Yoder AD, Goodman SM. A deep divergence time between sister species of Eidolon (Pteropodidae) with evidence for widespread panmixia. Acta Chiropterologica. 2014 Dec;16(2):279–92.

54. Goodman SM, Chan L, Nowak M, Yoder AD. Phylogeny and biogeography of western Indian Ocean Rousettus (Chiroptera:Pteropodidae). Journal of Mammalogy. 2010;91(3):593–606.

55. Almeida FC, Giannini NP, Simmons NB, Helgen KM. Each flying fox on its own branch: a phylogenetic tree for Pteropus and related genera (Chiroptera: Pteropodidae). Molecular Phylogenetics and Evolution [Internet]. 2014 Mar [cited 2014 Mar 23];(March). Available from: http://linkinghub.elsevier.com/retrieve/pii/S1055790314001092

56. Reynes J-M, Andriamandimby SF, Razafitrimo GM, Razainirina J, Jeanmaire EM, Bourhy H, et al. Laboratory surveillance of rabies in humans, domestic animals, and bats in Madagascar from 2005 to 2010. Advances in preventive medicine. 2011 Jan;2011:727821.

57. Brook CE, Ranaivoson HC, Broder CC, Cunningham AA, Héraud J-M, Peel AJ, et al. Disentangling serology to elucidate henipa- and filovirus transmission in Madagascar fruit bats. Journal of Animal Ecology [Internet]. 2019 Jul 1;88(7):1001–16. Available from: https://doi.org/10.1111/1365-2656.12985

58. Han Y, Du J, Su H, Zhang J, Zhu G, Zhang S, et al. Identification of diverse bat Alphacoronaviruses and Betacoronaviruses in China provides new insights into the evolution and origin of coronavirus-related diseases. Frontiers in Microbiology [Internet]. 2019;10:1900. Available from: https://www.frontiersin.org/article/10.3389/fmicb.2019.01900

59. Lim XF, Lee CB, Pascoe SM, How CB, Chan S, Tan JH, et al. Detection and characterization of a novel bat-borne coronavirus in Singapore using multiple molecular approaches. Journal of General Virology. 2019;100(10):1363–74.

60. Yinda CK, Ghogomu SM, Conceição-Neto N, Beller L, Deboutte W, Vanhulle E, et al. Cameroonian fruit bats harbor divergent viruses, including rotavirus H, bastroviruses, and picobirnaviruses using an alternative genetic code. Virus Evolution. 2018;4(1):1–15.

61. Huang C, Liu WJ, Xu W, Jin T, Zhao Y, Song J, et al. A bat-derived putative cross-family recombinant coronavirus with a reovirus gene. PLOS Pathogens [Internet]. 2016 Sep 27;12(9):e1005883.. Available from: https://doi.org/10.1371/journal.ppat.1005883

62. Obameso JO, Li H, Jia H, Han M, Zhu S, Huang C, et al. The persistent prevalence and evolution of cross-family recombinant coronavirus GCCDC1 among a bat population:a two-year follow-up. 2017;60(12):1357–63.

63. Goodman SM. Les chauves-souris de Madagascar [in French]. Antananarivo, Madagascar: Association Vahatra; 2011.

64. Golden CD, Bonds MH, Brashares JS, Rodolph Rasolofoniaina BJ, Kremen C. Economic valuation of subsistence harvest of wildlife in Madagascar. Conservation Biology. 2014 Jan 9;1–10.

65. Randrianandrianina F, Andriafidison D, Amyot F, Ramilijaona O, Ratrimomanarivo F, Racey PA, et al. Habitat use and conservation of bats in rainforest and adjacent human-modified habitats in eastern Madagascar. Acta Chiropterologica. 2006;8(2):429–37.

66. Cardiff SG, Ratrimomanarivo FH, Goodman SM. The effect of tourist visits on the behavior of Rousettus madagascariensis (Chiroptera: Pteropodidae) in the caves of Ankarana, northern Madagascar. Acta Chiropterologica. 2012 Dec;14(2):479–90.

67. Razafindrakoto N, Harwell A, Jenkins R. Bats roosting in public buildings: A preliminary assessment from Moramanga, eastern Madagascar. Madagascar Conservation & Development. 2011 Jan 11;5(2).

68. Jenkins RKB, Racey PA. Bats as bushmeat in Madagascar. Madagascar Conservation and Development. 2008;3(1):22–30.

69. Razanajatovo NH, Richard V, Hoffmann J, Reynes J, Razafitrimo M, Randremanana RV, et al. Viral etiology of Influenza-like illnesses in Antananarivo, Madagascar, July 2008 to June 2009. 2011;6(3).

70. Razanajatovo NH, Guillebaud J, Harimanana A, Rajatonirina S, Ratsima EH, Andrianirina ZZ, et al. Epidemiology of severe acute respiratory infections from hospital-based surveillance in Madagascar, November 2010 to July 2013. PLoS ONE. 2018;(July 2013):1–17.

71. Randremanana R, Andriamandimby S, Rakotondramanga JM, Razanajatovo N, Mangahasimbola R, Randriambolamanantsoa T, et al. The COVID-19 Epidemic in Madagascar: clinical description and laboratory results of the first wave, March-September 2020. Influenza and Other Respiratory Viruses. 2021;00:1–12.

72. Ranaivoson HC, Héraud J-M, Goethert HK, Telford SR, Rabetafika L, Brook CE. Babesial infection in the Madagascan flying fox, Pteropus rufus É. Geoffroy, 1803. Parasites & Vectors. 2019;12(1):51.

73. Brook CE, Bai Y, Dobson AP, Osikowicz LM, Ranaivoson HC, Zhu Q, et al. Bartonella spp. in fruit bats and blood-feeding ectoparasites in Madagascar. PLOS Neglected Tropical Diseases. 2015 Feb 23;9(2):e0003532..

74. Kalantar KL, Carvalho T, de Bourcy CFA, Dimitrov B, Dingle G, Egger R, et al. IDseq-An open source cloud-based pipeline and analysis service for metagenomic pathogen detection and monitoring. GigaScience. 2021;9(10):1–14.

75. Altschul SF, Gish W, Miller W, Myers EW, Lipman DJ. Basic local alignment search tool. Journal of Molecular Biology. 1990;215(3):403–10.

76. Kuraku S, Zmasek CM, Nishimura O, Katoh K. aLeaves facilitates on-demand exploration of metazoan gene family trees on MAFFT sequence alignment server with enhanced interactivity. Nucleic acids research. 2013;41(Web Server issue):22–8.

77. Katoh K, Rozewicki J, Yamada KD. MAFFT online service: Multiple sequence alignment, interactive sequence choice and visualization. Briefings in Bioinformatics. 2018;20(4):1160–6.

78. Darriba Di, Posada D, Kozlov AM, Stamatakis A, Morel B, Flouri T. ModelTest-NG: A new and scalable tool for the selection of DNA and protein evolutionary models. Molecular Biology and Evolution. 2020;37(1):291–4.

79. Kozlov AM, Darriba D, Flouri T, Morel B, Stamatakis A. RAxML-NG: A fast, scalable and user-friendly tool for maximum likelihood phylogenetic inference. Bioinformatics. 2019;35(21):4453–5.

80. Felsenstein J. Confidence limits on phylogenies: An approach using the bootstrap. Evolution. 1985;39(4):783–91.

81. Pattengale ND, Alipour M, Bininda-Emonds ORP, Moret BME, Stamatakis A. How many bootstrap replicates are necessary? Journal of Computational Biology. 2010;17(3):337–54.

82. Drummond AJ, Rambaut A, Shapiro B, Pybus OG. Bayesian coalescent inference of past population dynamics from molecular sequences. Molecular Biology and Evolution. 2005 Feb 9;22(5).

83. Bouckaert R, Vaughan TG, Barido-Sottani J, Duchêne S, Fourment M, Gavryushkina A, et al. BEAST 2.5: An advanced software platform for Bayesian evolutionary analysis. PLOS Computational Biology. 2019 Apr 8;15(4).

84. Rambaut A, Drummond AJ, Xie D, Baele G, Suchard MA. Posterior summarization in Bayesian phylogenetics using Tracer 1.7. Systematic Biology. 2018 Sep 1;67(5).

85. Drummond AJ, Rambaut A. BEAST: Bayesian evolutionary analysis by sampling trees. BMC Evolutionary Biology. 2007;7(1).

86. Yu G, Smith DK, Zhu H, Guan Y, Lam TTY. Ggtree: an R Package for visualization and annotation of phylogenetic trees with their covariates and other associated data. Methods in Ecology and Evolution. 2017;8(1):28–36.

87. Davies J. pySimPlot [Internet]. GitHub. [cited 2021 Sep 6]. Available from: https://github.com/jonathanrd/pySimPlot

88. Wu Y, Zhu X, Li N, Chen T, Yang M, Yao M, et al. CMRF-35–like molecule 3 preferentially promotes TLR9-triggered proinflammatory cytokine production in macrophages by enhancing TNF receptor-associated factor 6 ubiquitination. The Journal of Immunology. 2011 Nov 1;187(9).

89. Xu J, Hu J, Wang J, Han Y, Hu Y, Wen J, et al. Genome organization of the SARS-CoV. Genomics, Proteomics & Bioinformatics. 2003;1(3):226–35.

90. Kim D, Lee JY, Yang JS, Kim JW, Kim VN, Chang H. The architecture of SARS-CoV-2 transcriptome. Cell. 2020;181(4):914-921.e10.

91. Li F. Receptor recognition and cross-species infections of SARS coronavirus. Antiviral Research. 2013;100(1):246–54.

92. Tao Y, Shi M, Chommanard C, Queen K, Zhang J, Markotter W, et al. Surveillance of bat coronaviruses in Kenya identifies relatives of human coronaviruses NL63 and 229E and their recombination history. Virology. 2017;91(5):1–16.

93. Halpin K, Rota P. A review of hendra virus and nipah virus Infections in man and other animals. Sing A, editor. Zoonoses - Infections Affecting Humans and Animals: Focus on Public Health Aspects [Internet]. 2014 Aug 22;997–1012. Available from: https://www.ncbi.nlm.nih.gov/pmc/articles/PMC7120151/

94. Wells HL, Letko M, Lasso G, Ssebide B, Nziza J, Byarugaba DK, et al. The evolutionary history of ACE2 usage within the coronavirus subgenus sarbecovirus. Virus Evolution [Internet]. 2021 Jan 20;7(1). Available from: https://doi.org/10.1093/ve/veab007

95. Andrianaivoarivelo RA, Ramilijaona OR, Racey PA, Razafindrakoto N, Jenkins RKB. Feeding ecology, habitat use and reproduction of Rousettus madagascariensis Grandidier, 1928 (Chiroptera: Pteropodidae) in eastern Madagascar: 2011;75(1):69–78. Available from: https://doi.org/10.1515/mamm.2010.071

96. Rasambainarivo F, Ramiadantsoa T, Raherinandrasana A, Randrianarisoa S, Rice BL, Evans M v, et al. Prioritizing COVID-19 vaccination efforts and dose allocation within Madagascar. medRxiv [Internet]. 2021 Jan 1;2021.08.23.21262463. Available from: http://medrxiv.org/content/early/2021/08/25/2021.08.23.21262463.abstract

97. Rabalski L, Kosinski M, Mazur-Panasiuk N, Szewczyk B, Bienkowska-Szewczyk K, Kant R, et al. Zoonotic spillover of SARS-CoV-2: mink-adapted virus in humans. bioRxiv [Internet]. 2021 Jan 1;2021.03.05.433713. Available from: http://biorxiv.org/content/early/2021/03/05/2021.03.05.433713.abstract

98. Wacharapluesadee S, Duengkae P, Chaiyes A, Kaewpom T, Rodpan A, Yingsakmongkon S, et al. Longitudinal study of age-specific pattern of coronavirus infection in Lyle’s flying fox (Pteropus lylei) in Thailand. Virology Journal [Internet]. 2018;15(1):38. Available from: https://doi.org/10.1186/s12985-018-0950-6

99. Annan A, Baldwin HJ, Corman VM, Klose SM, Owusu M, Nkrumah EE, et al. Human betacoronavirus 2c EMC/2012-related viruses in bats, Ghana and Europe. Emerging infectious diseases [Internet]. 2013 Mar;19(3):456–9. Available from: https://pubmed.ncbi.nlm.nih.gov/23622767

100. Brook CE, Ranaivoson HC, Andriafidison D, Ralisata M, Razafimanahaka J, Héraud J-M, et al. Population trends for two Malagasy fruit bats. Biological Conservation [Internet]. 2019;234:165–71. Available from: https://www.sciencedirect.com/science/article/pii/S0006320718316744

101. Weizhong L. cdhit [Internet]. GitHub. [cited 2021 Sep 6]. Available from: https://github.com/weizhongli/cdhit

